# Macaque dorsal premotor cortex exhibits decision-related activity only when specific stimulus-response associations are known

**DOI:** 10.1101/412528

**Authors:** Megan Wang, Christéva Montanède, Chandramouli Chandrasekaran, Diogo Peixoto, Krishna V. Shenoy, John F. Kalaska

**Author notes:** These first authors contributed equally (Megan Wang, Christéva Montanède). These senior authors contributed equally (Krishna V. Shenoy, John F. Kalaska). Correspondence (M.W.) and (J.F.K.).

## Abstract

How deliberation on sensory cues and action selection interact in decision-related brain areas is still not well understood. Here, monkeys reached to one of two targets, whose colors alternated randomly between trials, by discriminating the dominant color of a checkerboard cue composed of different numbers of squares of the two target colors in different trials. In a “Targets First” task the colored targets appeared first, followed by the checkerboard; in a “Checkerboard First” task, this order was reversed. After both cues appeared in both tasks, responses of dorsal premotor cortex (PMd) neurons covaried with action choices, strength of evidence for action choices, and RTs--- hallmarks of decision-related activity. However, very few neurons were modulated by checkerboard color composition or the color of the chosen target, even in the Checkerboard First task. These findings implicate PMd in the action-selection but not the perceptual components of the decision-making process in these tasks.

## Introduction

A fundamental role of the brain is to guide the physical interactions of the individual with her environment. This requires continual decisions about action choices by deliberating upon sensory evidence from the world to decide in favor of one choice over other alternative actions (Brody and Hanks, 2016; Cisek and Kalaska, 2010; Freedman and Assad, 2011; Gold and Shadlen, 2007; Romo and de la Fuente, 2013; Shadlen and Kiani, 2013). In some situations, the evaluation of the sensory evidence can occur conjointly with decisions about action choices, for instance while trying on different pairs of boots at a shoe store before deciding on which pair to buy. In others, assessment of the evidence and selection of the actions that implement the decision can be dissociated in time, for instance by choosing which pair of boots to buy on the store’s web site before going to the store to purchase them. Of course, we can also make an abstract categorical decision about whether or not we like a particular pair of boots even when we have no intention to buy and wear them.

A large literature implicates several premotor brain areas associated with operantly-conditioned motor responses in sensorimotor decision-making processes leading to action choices (Brody and Hanks, 2016; de Lafuente et al., 2015; Glimcher, 2001; Huda et al., 2018; Roitman and Shadlen, 2002; Romo et al., 2004; Shadlen and Newsome, 1996). In an extensive series of studies in the oculomotor system, subjects chose between saccade targets in known locations by estimating the net direction of visual motion in random-dot kinematogram (RDK) stimuli with variable amounts of coherent motion (Britten et al., 1992). Neural activity in several cortical and subcortical saccade-related premotor structures showed putative correlates of the sensorimotor deliberation process leading to a saccade (Ding and Gold, 2010, 2012; Horwitz and Newsome, 2001; Kim and Shadlen, 1999; Roitman and Shadlen, 2002; Shadlen and Newsome, 1996, 2001).

Similarly, in the arm motor system, the dorsal premotor cortex (PMd) is a strong candidate region in which sensory instructional and action-related information may interact to guide voluntary arm movements (Afshar et al., 2011; Boussaoud and Wise, 1993; Chandrasekaran et al., 2017; Churchland and Shenoy, 2007; Cisek, 2007; Cisek and Kalaska, 2010; Crammond and Kalaska, 1996, 2000; di Pellegrino and Wise, 1991, 1993; Hoshi and Tanji, 2000, 2006; Kalaska and Crammond, 1995; Kaufman et al., 2015; Messier and Kalaska, 2000; Mitz et al., 1991; Riehle and Requin, 1989; Thura and Cisek, 2014; Wise et al., 1997). Those studies showed that PMd cells respond to sensory instructional cues that provide full or partial information about intended actions, and also signal the final action choices made by the subjects. For instance, reach-related neurons in PMd can first generate representations of two potential reach choices and then the final selected target when monkeys choose between two color-coded targets according to monochromatic color cues (Cisek and Kalaska 2005). PMd neurons can express learned stimulus-response associations while monkeys observe the task-related sensory events on a monitor but do not actively perform the task (Cisek and Kalaska, 2004). Finally, PMd activity can covary with higher-level abstract action-related concepts before the action choices are fully specified, such as the general goal of a future action (Nakayama et al., 2008) or a visuomotor task rule (GO/NOGO; Wallis and Miller, 2003).

An ongoing challenge is to determine to what degree the evolving decision-related activity in premotor circuits reflects a process of deliberation on the sensory evidence associated with the perceptual decision or with the choice of action. Indeed, it is still an open question whether distinct perceptual versus motor decision processes can be disentangled in premotor circuits or whether are they inseparably linked in a broader “embodied” sensorimotor intentional process in which the overall behavioral context and goals determine both the appropriate set of action options and the salient sensory inputs that guide action selection (Churchland et al., 2011; Gold and Shadlen, 2007; Cisek and Kalaska, 2010; Shushruth et al., 2018; Huda et al., 2018). With rare exceptions (e.g. Bennur and Gold, 2011; Gold and Shadlen, 2003; Horwitz et al., 2004), in the vast majority of studies of perceptual decision-making in RDK tasks, the subjects knew how sensory evidence would be mapped onto saccade-target choices before the salient sensory input began. As a result, the subjects could select and prepare the “report” motor response simultaneously with the sensory evidence acquisition process that informed the motor decision. In addition, the mapping of coherent-motion sensory evidence onto action choice was usually along the same directional axis and thus not dissociable, making it difficult to disambiguate to what degree the premotor neural responses reflected sensory-evidence or motor-decision processes.

Our labs have been studying the role of PMd in reach decisions in tasks in which subjects must discriminate the dominant color of a multi-colored checkerboard “decision cue” to select between two color-coded targets. We controlled the amount of sensory evidence supporting the action decision by varying the relative numbers of squares of the two target colors in the checkerboard between trials (its color evidence “coherence” and dominant color; see Methods for the definition of “coherence” in this task context), analogous to the variable motion coherence and net motion direction of RDK stimuli. In our initial studies, we used a Targets First (TF) task in which the action choices (colored target locations) were known in each trial before the checkerboard appeared (Chandrasekaran et al., 2017; Coallier and Kalaska, 2014; Coallier et al., 2015). We both found that neural activity in PMd of monkeys was correlated with the strength of checkerboard evidence supporting a reach target and with the direction of the reach target the monkeys chose (Chandrasekaran et al., 2017; Coallier et al., 2015; Peixoto et al., 2018, bioRxiv doi: 10.1101/283960). These neural data are consistent with an intentional framework for decision-making (Cisek and Kalaska 2010; Shadlen et al., 2008) in which sensory evidence-related activity is expressed in brain areas that generate the motor report of the decision (Roitman and Shadlen, 2002; Romo et al., 2004). Nevertheless, the TF task shares the same interpretational limitation of many prior studies that the onset of the checkerboard can initiate the perceptual-deliberation and action-selection processes simultaneously.

To assess whether and when PMd activity expresses sensory- or action-related components of the sensorimotor decision process, we both implemented a task variant, the Checkerboard First (CF) or Checkerboard First with Delay (CFD) task, in which the sensory decision cue is presented before the associated action choices are revealed (c.f. Bennur and Gold, 2011; Gold and Shadlen, 2003; Horwitz et al 2004). In theory, this could permit subjects to make a categorical perceptual decision about the dominant checkerboard color before the colored targets appear. Human subjects showed behavioral evidence that they made a non-intentional categorical perceptual decision in a CF task (Coallier and Kalaska, 2014). Their reaction times (RTs) were much shorter and far less dependent on checkerboard coherence than in a TF task. This suggested that they decided on the dominant color of the checkerboard while observing it before the colored target locations were known and then used that prior perceptual decision to choose the color-matching reach target rapidly after they appeared.

We recorded neural activity in PMd while monkeys performed the TF and CF (monkey Z) or TF and CFD (monkey T) tasks. We sought PMd neural correlates of decision-making processes, defined operationally here as differential neural activity that predicted any aspect of the monkeys’ task-related decisions, including the spatial location or direction of the chosen target, the dominant color of the checkerboard or the color of the chosen target. At the same time, we acknowledge that these data are correlational and are not proof of a causal relationship between PMd activity and perceptual or motor decisions. A key hypothesis was that if PMd reflects perceptual decision-making information independent of full prior knowledge of associated action choices, then neural activity during the checkerboard observation epoch of the CF and CFD tasks would reflect its dominant color, the critical property of the checkerboard that determines the monkeys’ action decisions. We confirmed that our data in the TF task were consistent with previous findings (Chandrasekaran et al., 2017; Coallier et al., 2015); neural activity following checkerboard onset in both monkeys reflected the level of checkerboard coherence favoring a target direction, the action choices and RTs, but not the checkerboard’s dominant color or the color of the chosen target. Monkeys T and Z showed different RT trends and different degrees of responsiveness of PMd neurons to the appearance of the checkerboard in the CF/CFD tasks. Most importantly, however, virtually no PMd neuron in either monkey showed a differential response to the checkerboard’s dominant color, either before or after the targets appeared in the CF/CFD tasks, or to the color of the chosen target. Our results indicate that PMd does not express correlates of the critical color perceptual decision process that informs the reach target choices in either the TF or CF/CFD tasks. Instead, PMd neurons become differentially active only when complete information about the stimulus-response associations that determine action choices is available, and their activity primarily reflects the properties of those actions (e.g., reach direction), the strength of evidence supporting those actions, and the temporal dynamics of the action decisions.

## Results

### A task design to dissociate perceptual decisions from action-selection decisions in time

Two rhesus macaque monkeys performed variants of a sensorimotor decision-making task (Figure 1A). Their goal was to determine the dominant color of a multi-colored checkerboard, and to report that color by reaching to the correspondingly colored target. The two targets were always presented in the same two opposite spatial locations along the estimated preferred movement axis for each unit for hundreds of successive trials during recording sessions for monkey Z; monkey T’s target locations were permanently fixed to the left and right of the central start position. As a result, both monkeys could potentially anticipate the spatial locations of the targets at the start of each trial. However, target colors were assigned randomly on each trial, so the monkeys did not know which color would be assigned to which target until the targets appeared.

**Figure 1.**
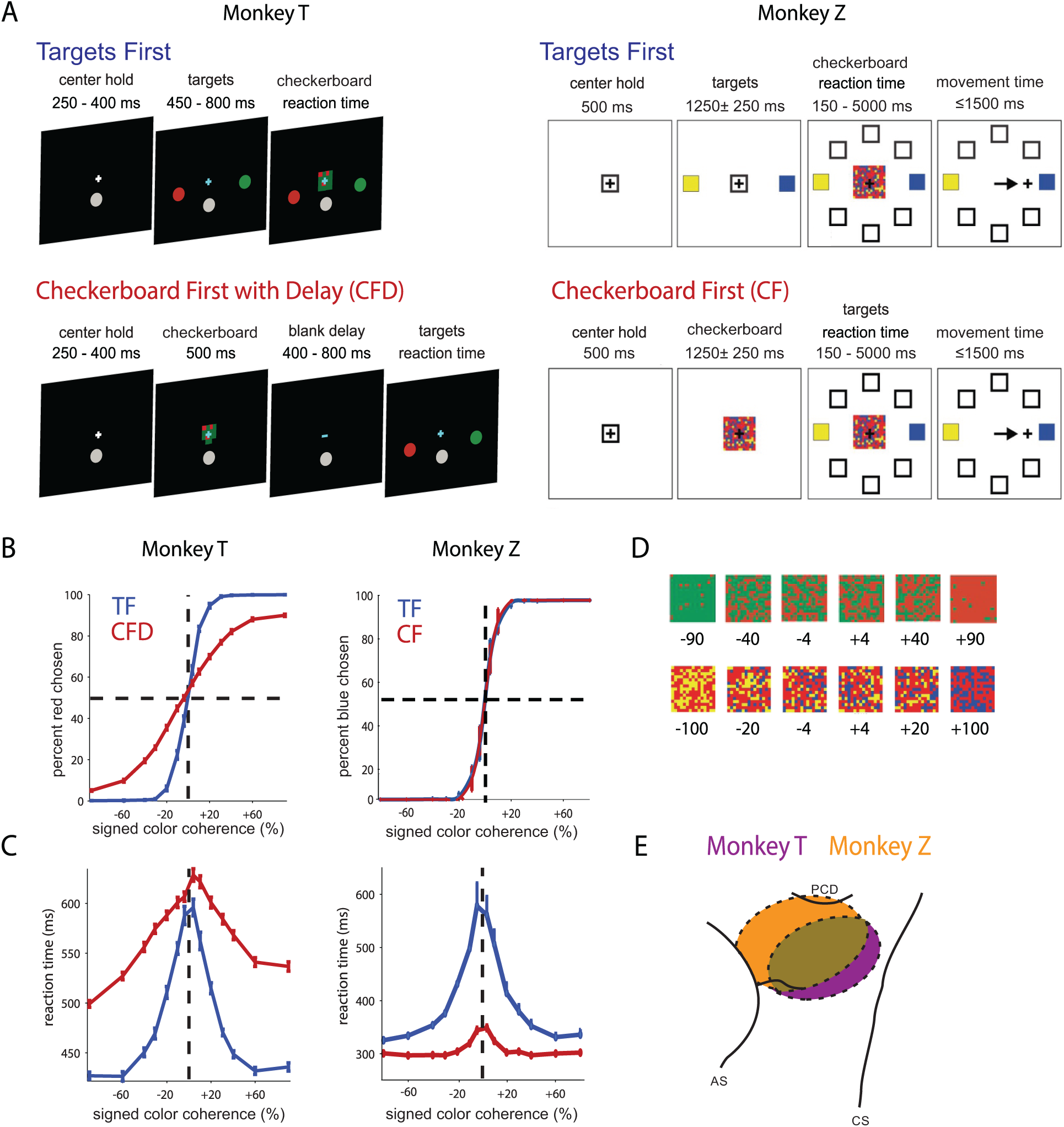
Choice behavior of two monkeys in Target First (TF) and Checkerboard First (CF) tasks. **A)** Two monkeys from two labs performed variants of a checkerboard color discrimination task. The TF task design (top row) follows the usual convention of presenting possible action choices – here, colored targets – before presenting a decision cue – here, a colored checkerboard. The goal is to report the dominant color of the checkerboard by reaching to the corresponding colored target. Note that in this design, perceptual decision-making (checkerboard color discrimination) and action selection (choosing the target with the matching color) both occur in the reaction-time interval between the appearance of the checkerboard and the onset of the reaching movement and in theory can happen simultaneously. In the CF and CF with Delay (CFD) task design (bottom row), the order of presentation is reversed to attempt to dissociate perceptual decision-making and action selection in time. The colors of the targets are assigned randomly to the targets on each trial. **B)** Psychometric curves of the probability of Red or Blue target color choices (mean ± SEM) as a function of signed checkerboard color coherence (orange curves: CFD task for monkey T, and CF task for monkey Z; blue curves: TF tasks). Signed color coherence is the difference in number of colored squares for each category (e.g. # red squares - # green squares) divided by the total number of task-relevant squares in the checkerboard. Positive values signify predominantly R or B checkerboards, and negative values signify predominantly G or Y checkerboards. As the strength of color evidence in the checkerboards supporting a reach to an R/B target increased, the probability of reaches to those targets increased. Psychometric curves of the probability of G or Y target color choices are mirror-symmetrical. **C)** Chronometric curves of reaction times (RTs; mean and SEM) as a function of signed checkerboard color coherence. As the strength of color evidence in the checkerboard supporting a reach to one colored target over the other increased, RT durations tended to decrease. See also Supplemental Figure 1. **D)** Examples of R/G checkerboards for monkey T (top row) and B/Y checkerboards for monkey Z (bottom row), labeled with their level of coherence (units are %). **E)** Recordings were performed in dorsal premotor cortex (PMd) in the hemisphere contralateral to the reaching arm. Data include units recorded from single electrodes and linear arrays. This schematic map illustrates the approximate locations of recording sites based on stereotactic coordinates. Histology has not yet been done on either monkey. PCD, precentral dimple; AS, arcuate sulcus; CS, central sulcus.

In this study, the critical sensory evidence is provided by a multi-colored checkerboard stimulus that presents different numbers of colored squares supporting the two reach choices, somewhat analogous to the coherent motion strength in RDK stimuli. A key differentiator for this study is that the color of an object does not have any inherent association with any parameter of a reach movement, such as target spatial location or reach direction. Color only becomes action-relevant in our tasks by application of an arbitrary stimulus-response rule; the subjects decide on the dominant color of the checkerboard and use a color-location matching rule to associate it with the target of the same color.

The Targets First (TF) task variant followed the event timeline used in many sensorimotor decision tasks, in which the colored targets appeared before the checkerboard decision cue (Chandrasekaran et al., 2017; Coallier and Kalaska 2014; Coallier et al., 2015; Kim and Shadlen, 1999; Shadlen and Newsome, 2001). As a result, deliberation about dominant checkerboard color could occur concomitantly with planning for the reach, because each color was already associated with a specific target location. Crucially, in the Checkerboard First (CF) and Checkerboard First with Delay (CFD) tasks, the order of the two sensory events was reversed. The checkerboard appeared first, and the monkeys could in theory deliberate upon the checkerboard’s dominant color but could not prepare a specific motor response to report it because the colored targets had not yet appeared. The monkeys were free to initiate a reach to a target at the time of their choosing after the second visual cue appeared in each task. The details of the task event sequence and timing varied between the two laboratories (Figure 1A; see also Methods).

### Task performance of the two monkeys was similar in the TF task but different in the CF/CFD tasks

Figure 1B, C (left) show monkey T’s task performance during all neural recording sessions. To facilitate comparison of task performance, monkey Z was tested in separate behavioral sessions without neural recordings, using 7 checkerboard coherences ranging from 4% – 80% (Figure 1B, C, right). A reduced set of checkerboard color coherences (4%, 20% and 100%) was used during neural recordings with monkey Z; task performance was completely consistent with the trends described here (Table 1).

**Table 1.** Mean success rate (%) and mean RT ± s.e.m. (ms) for the data files collected during all neural recording sessions in monkey Z.

| Checkerboard coherence | 4% | 20% | 100% |
| --- | --- | --- | --- |
| **TF task** |  |  |  |
| Success rate | 66.1% | 95.9% | 100.0% |
| Mean RT ± s.e.m. (ms) | 529.1±18.2 | 426.3±11.6 | 323.7±4.5 |
| **CF Task** |  |  |  |
| Success rate | 67.2% | 97.3% | 100.0% |
| Mean RT ± s.e.m. (ms) | 341.5±8.1 | 303.5±6.3 | 298.2±5.6 |

The performance of both monkeys was strikingly similar in the TF task. Both performed at above chance levels with 4% checkerboards (T: 63.0% correct target choices; Z: 71.6%) and reached an asymptote at ∼100% success rates with checkerboard coherences of 20-90% (Figure 1B, blue curves). The psychometric curves were symmetric and centered on 0% color coherence, indicating that neither monkey showed a major bias towards one color over the other. The reaction times (RTs) were maximal for the 4% checkerboards (T: 585±6 ms (s.e.m.), Z: 583±36 ms) and decreased systematically as checkerboard coherence increased, approaching an asymptote at minimal values at 60-90% checkerboards (T: 435±3 ms for 90% coherence, Z: 333±9 ms for 80% coherence; Figure 1C, blue curves). Thus, the checkerboard color coherence had effects on the choice behavior and reaction times that were very reminiscent of the effect of motion direction coherence in RDK tasks (Ding and Gold, 2012; Palmer et al., 2005; Roitman and Shadlen, 2002), despite the many differences in the stimuli. Indeed, the RT distributions across all checkerboard coherences could be readily fit by a standard drift-diffusion model to a fixed decision bound for each monkey (data not shown).

The performance of the two animals differed in the CF/CFD task. Monkey T did not achieve 100% performance in the CFD task even at the strongest color coherence (lapse rate 7.4% for 90% color coherence), and the psychometric curve had a lower slope in the CFD task compared to the TF task (Figure 1B left, red curve compared to blue curve). In contrast, the psychometric curve for monkey Z in the CF task was essentially identical to that in the TF task (Figure 1B, right, red curve), possibly in part because of the longer checkerboard observation period (1000-1500 ms) in the CF task than the CFD task and its continued presence until the monkey initiated a reach.

To compare psychophysical thresholds for checkerboard color coherence between the two tasks, we fit the “folded” psychometric curves (% correct as a function of unsigned coherence) to cumulative Weibull functions (Roitman and Shadlen, 2002). For monkey T, the psychophysical threshold was 10.4% in the TF task, and increased to 41.9% in the CFD task. For monkey Z, psychophysical thresholds were lower (TF task: 7.0%; CF task: 6.8%) than for monkey T, and very similar between tasks.

We also observed differences in the RT trends in the CF/CFD tasks between the two monkeys. Monkey T’s RTs were systematically slower in the CFD task than in the TF task at all checkerboard coherences. This was most pronounced for the checkerboards with the strongest color coherence, even though monkey T only performed those trials at >92% success rates compared to ∼100% in the TF task (Figure 1C, left, red line). In contrast, monkey Z’s RTs were shorter in the CF task than the TF task at all coherences, with the largest reduction for checkerboards with the lowest coherences (Figure 1C, right, red line). There was a much smaller dependence on checkerboard coherence in the CF task; RTs for 4% checkerboards (350±13 ms) were only ∼45 ms longer than for the 80% checkerboards (303±7 ms). Furthermore, Monkey Z’s RTs for the 4% checkerboards in the CF task were only modestly longer than for the 80% checkerboards in the TF task, which were in turn ∼30ms longer than for the 80% checkerboards in the CF task.

Monkey Z was re-trained and tested in a modified CF task with identical temporal structure to the CFD task (see Methods). Like monkey T, monkey Z showed a decrease in success rates for the checkerboards with stronger color coherence in the CFD task, with a lapse rate of 10.5% (Supplemental Figure 1A). However, unlike monkey T, monkey Z continued to show nearly all the temporal savings in the CFD task that had been observed in the CF task. Its RTs for the 4% checkerboards (348±8 ms) were essentially identical to that in the CF task, and were only slightly prolonged for the 80% checkerboards (328±7 ms) compared to the CF task (Supplemental Figure 1B).

To summarize, both monkeys performed the tasks with psychometric and chronometric trends that strongly reflected the strength of color evidence in the checkerboards. The near-perfect success rates with 20%-90% color-coherence checkerboards in the TF task showed that the monkeys successfully interpreted the color evidence in the checkerboards and correctly chose the target with the same color, regardless of its spatial location. Their ability to choose the correctly-colored target diminished rapidly as the checkerboard color coherence decreased below 20%. Monkey T’s RTs were prolonged rather than reduced in the CFD task, but its choice behavior showed that it retained some unknown but task-salient information about the checkerboard during the memory-delay period and used that stored information to make target choices whose success rates increased and RTs decreased systematically with the evidence strength after the colored targets appeared. Monkey Z showed a marked reduction in RTs in the CF and CFD tasks that was largest for checkerboards with the weakest color coherence, as did human subjects (Coallier and Kalaska 2014). While alternative explanations are possible, we suggest that the shorter RTs in the CF/CFD tasks are a behavioral sign that monkey Z made a categorical perceptual decision about the dominant color of the checkerboards during the checkerboard observation period, resulting in a substantial shortening of the RTs after the colored targets appeared.

These behavioral findings indicated that both monkeys made use of the sensory information available during the checkerboard observation period of the CF/CFD tasks, but in different ways that may have resulted in different degrees of temporal separation of the perceptual-deliberation and action-selection aspects of the tasks. We next asked how and when neural correlates of the perceptual and action decisions were expressed in PMd as a function of the amount of information available about specific action choices. Strong correlates with the color of the chosen target or with the amount of color evidence in the checkerboard independent of action choices would indicate that PMd expresses activity reflecting the process of perceptual deliberation on the task-salient sensory evidence. Neural correlates with the direction of the chosen target independent of the color evidence support a role in action selection, and neural correlates with the amount of non-color directional evidence favoring an action choice could support a role in acquiring sensory evidence supporting specific action decisions.

### Recordings in PMd: unit selection criteria

We recorded neural data from PMd in the hemisphere contralateral to the performing arm, including the left hemisphere of monkey T using standard single microelectrodes or single multi-port linear-array electrodes, and both hemispheres in monkey Z using standard single microelectrodes (Figure 1E). We recorded from a total of 499 units in monkey T during the CFD task, of which a subset of 351 units was also recorded during the TF task. For monkey T, the two target locations were always to the left and right of the starting hand position, and units were selected for analysis if they responded during any epoch of the task. All recorded neurons in monkey Z were pre-screened for task-related responses using 8-direction “1-Target” and “2-Target” instructed-delay tasks described previously (Cisek and Kalaska 2005; Coallier et al 2015). Complete data sets were then collected from 104 isolated units in both the TF and CF tasks (41 and 63 units from the left and right hemispheres respectively). During those recordings, the targets were placed in each cell’s task-related preferred movement direction and diametrically opposite direction as assessed in the 8-direction tasks.

### Neurons exhibited heterogeneous activity in response to the first sensory information provided in each task

The principal objective of the study was to assess the degree to which differential neural activity in PMd reflects perceptual-deliberation and action-selection processes. To begin, we briefly describe how the PMd neurons responded during observation of the first visual cue in each task, when only partial (TF task, color-coded target cues) or no (CFD/CF task, colored checkerboard) specific action-choice information is available.

The example unit from monkey T in Figure 2 (left) showed no change in activity during the first-cue observation period in either task, which continued into the memory-delay period of the CFD task. This was representative of the large majority of units in monkey T. The example unit from monkey Z (Figure 2, right) also showed no evident response during the Targets-observation period in the TF task, but exhibited a small fluctuation in activity 100-300ms after checkerboard appearance during the Checkerboard-observation period of the CF task. We did a bin-wise search to identify significant rapid changes in activity in response to the first cue (see Methods). Only 1/350 and 0/499 units in monkey T showed a detectible rapid activity change in response to the first cue in the TF and CFD tasks respectively. Many of the neurons in monkey Z showed rapid changes in activity shortly after the appearance of the first visual cue in the TF (42/104) and CF tasks (52/104); 35/42 and 44/52 of those responses were detected in the time period 100-300ms after the cues appeared (Figure 2, 3, Supplemental Figure 2). Subsequent analyses assessed whether PMd neurons in either monkey expressed differential decision-making activity that predicted the monkeys’ sensory or action decisions during the initial visual-cue observation period before the specific stimulus-response mapping was fully defined in each trial, or only after the second visual cue appeared.

**Figure 2.**
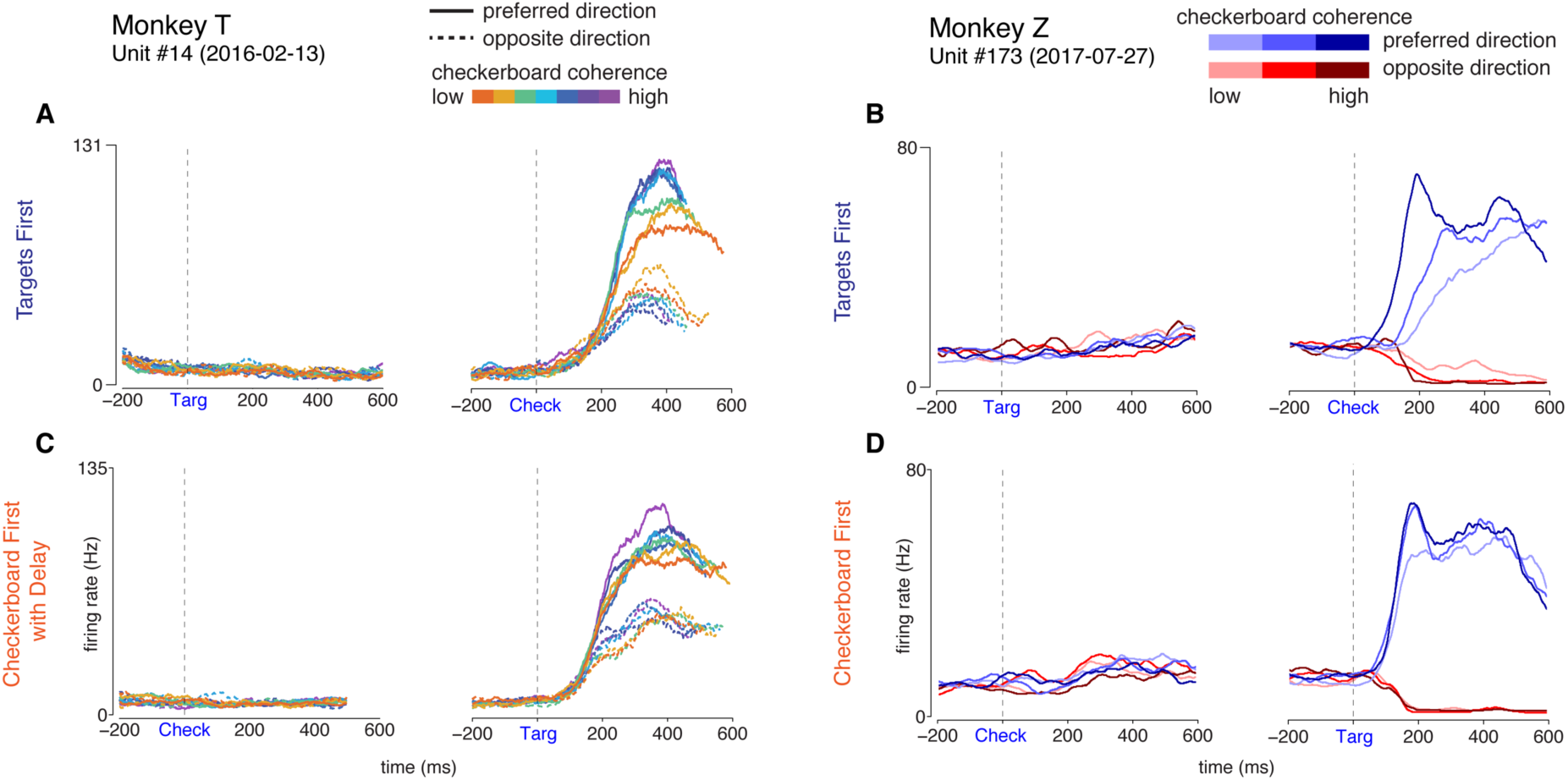
Example single-unit activity profiles in the TF and CF/CFD tasks. Data are aligned to the appearance of the first visual cue (left) and the second visual cue (right) in each task. **A,B)** TF Task. Consistent with previous work with similar tasks, the rate of change in neural activity after the appearance of the second visual cue (Checkerboard, “Check”)) in the TF task correlates both with checkerboard coherence and with reach direction. Neither neuron responded to the appearance of the first visual cue, the Targets (“Targ”), in that task. **C,D)** CFD/CF Task. In the CF and CFD tasks, the same units show a reduced effect of checkerboard color coherence on the rate of change of activity after the Targets appear. The unit from monkey T showed no change in activity during the initial Checkerboard-observation period of the CF task, but the neuron from monkey Z showed a small rapid response change, that was detected 280ms after Checkerboard appearance (see text).

**Figure 3.**
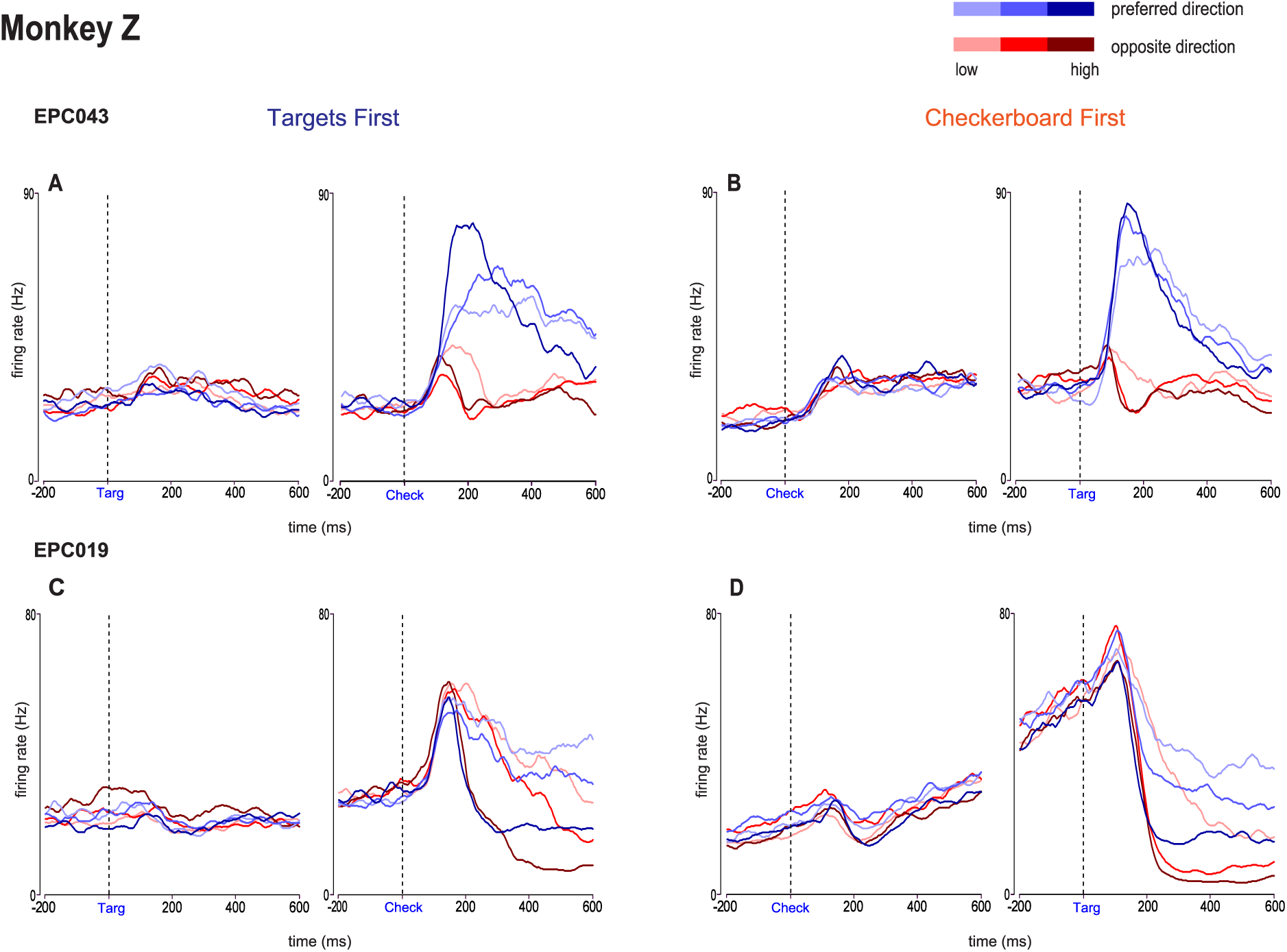
Examples of units recorded in monkey Z that responded to the first and second cues in the TF and CF tasks. See also Supplemental Figure 2. **A,B)** Unit EPC043 emitted a transient increase in activity detected at 160ms after the Targets appeared in the TF task (A), and a stronger sustained response beginning 140ms after the appearance of the Checkerboard in the CF task (B). **C,D)** Unit EPC019 emitted a small rise in activity that was detected at 240ms after Targets appearance in the TF task (C). In the CF task (D), there was a brief increase in activity detected at 140ms after the appearance of the Checkerboard, followed by a transient suppression and then a pronounced ramp increase for the remaining duration of the Checkerboard-observation period.

### Neural activity reflected checkerboard coherence and action choices only after the appearance of the second visual cue in both tasks

After the second visual cue appeared in each task, the monkeys had all the sensory information needed to complete the sensorimotor decision process and select a reach target. Unit activity in the TF task in both monkeys (Figure 2, 3) showed responses similar to that described previously, including differences in activity dependent on the direction of reach and on the coherence level of the checkerboards (Chandrasekaran et al., 2017; Coallier et al., 2015). In the CFD task, the example unit from monkey T had a smaller range of rates of change of discharge in both reach directions as a function of the no-longer visible checkerboard. The example units from monkey Z showed nearly identical rapid rates of change in activity for the 100% and 20% checkerboards and only a modestly slower rate of change for the 4% checkerboards (Figure 2, 3; Supplemental Figure 2). To quantify these responses, we estimated the slope of a differential directional “choice selectivity signal” associated with each checkerboard coherence during a time window 0-300 ms after the appearance of the two visual cues in the two tasks (Chandrasekaran et al., 2017; Meister and Huk, 2013; see Methods).

The slope of the choice selectivity signal 0-300 ms after the first cue appeared in each task yielded similar very low values in both tasks in each monkey (Figure 4 A,C); this indicated that the activity during this epoch did not predict the action choices of the monkeys, even while observing the color-coded targets in the TF task. No change in firing rate as a function of checkerboard coherence was seen in the TF task, as should be expected because the checkerboard had not yet appeared. In contrast, during the initial checkerboard-observation epoch in the CF/CFD tasks, color coherence information was present on the screen, but no information about how a color would be converted into a reach, and the mean choice selectivity slope also did not vary with coherence in either monkey.

**Figure 4.**
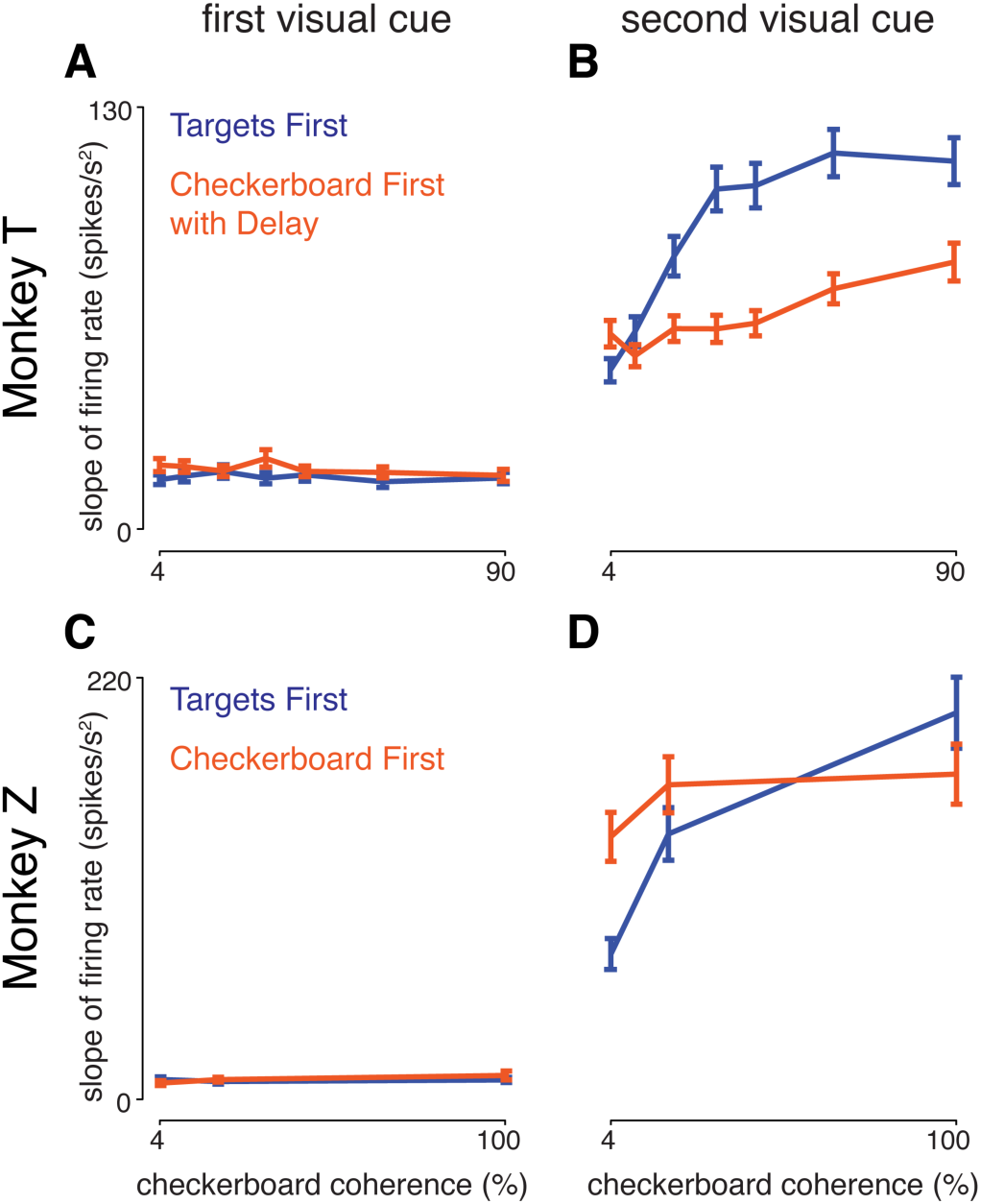
Neural activity reflected action choice and checkerboard coherence following the appearance of the second visual cue in each task. Plotted are the discharge rate slope (mean ± s.e.m.) of the choice selectivity signal as a function of checkerboard coherence. The choice selectivity signal is the difference in activity for trials in the two opposite reach directions (Chandrasekaran et al., 2017). For each unit, the choice selectivity signal recorded 0-300ms after the appearance of the first visual cue (**A,C**) and the second visual cue (**B,D**) was calculated separately for trials with each checkerboard coherence in the TF (blue) and CFD/CF tasks (orange), and then fit to a linear ramp function to estimate the short-latency rate of change of activity in response to the two visual cues (see Methods). While observing the first visual cue, choice selectivity signal slopes were small and did not vary as a function of checkerboard coherences, in both tasks in monkey T (**A**) and monkey Z (**C**), even while observing the checkerboards in the CFD/CF task. This indicated that the first cue did not evoke a directional signal in neural activity, as should be expected. Following the appearance of the second visual cue in each task however, strong choice selectivity signals appeared in PMd in both monkey T (**B**) and monkey Z (**D**). Slopes of the signals were strongly modulated by checkerboard coherence in the TF task (blue) but much less sensitive to checkerboard coherence in the CFD/CF task (orange) in both monkeys.

The slopes of choice selectivity signals after the second visual cue appeared were large in both monkeys and varied systematically with the level of checkerboard coherence, so that activity differentiating the two chosen reach directions increased more quickly as the coherence increased (Figure 4 B,D). This was more prominent in the TF task than the CF/CFD task. The increase in slope values in single units in the TF task was statistically significant between the 4% and 20% coherences (2-tailed paired t-test; Monkey T: p = 8.35E-11, Monkey Z: p = 1.05E-11), and between the 20% and highest-coherence checkerboards (Monkey T: p = 1.23E-06, Monkey Z: p = 6.59E-09). Slope values were higher in monkey Z than in monkey T, in part likely reflecting the sampling bias in monkey Z to maximize the directionality of recorded unit activity. In the CFD task, the slopes of choice selectivity signals of single units in monkey T increased at higher coherence (slopes at 20% vs 90%, p = 3.08E-04) but not at lower coherences (slopes at 4% vs 20%, p = 0.7302). In contrast, the slopes of the choice selectivity signals for monkey Z in the CF task increased significantly from the 4% to the 20% checkerboards (p = 1.45E-05), but were similar for the 20% and 100% checkerboards (p = 0.26). The slopes were generally lower in the CFD task than in the TF task in monkey T, especially at higher coherences (2-sample t-test comparing slopes across tasks; for 4%, p = 0.05; for 20%, p = 1.4E-03; for 90%, p = 7.83E-04). The slopes were significantly higher for monkey Z in the CF task than the TF task for the 4% checkerboards (p = 1.05E-05), non-significantly higher for the 20% checkerboards (p = 0.011) and significantly lower for the 100% checkerboards (p = 5.6E-04).

In summary, neural activity after the second visual cue reflected the direction of chosen reach in both tasks, including trials with both correct and incorrect target choices, and the rate of change of the choice selectivity signal was strongly dependent on checkerboard coherence in the TF task in both monkeys. The effect of checkerboard coherence on choice selectivity slope was reduced in the CFD task for monkey T, and was even less prominent in the CF task for monkey Z.

### Neural responses reflect chosen reach direction and color-independent evidence towards a reach direction, but not chosen target color or signed checkerboard color coherence

To better understand how the correlations of neural activity with different task variables evolved over time, we did a linear regression analysis (*regress*; Matlab 2018A, the Mathworks Inc) of the single-trial activity during sequential non-overlapping 20ms time bins during the trial. We aligned single-trial data to the onset of the first and second visual cues in both tasks. For each unit and each time point, we regressed the single-trial firing rates on the following predictor variables: chosen reach direction, chosen target color, signed color coherence (the amount of evidence for one color over the other), and signed directional coherence (the amount of color-independent evidence supporting a reach direction). The last predictor requires knowledge of the specific target location-color associations in each trial. We included both correct and incorrect choices so that the color and direction of the chosen target reflects the monkeys’ interpretation of the sensory evidence rather than the correct dominant color of the checkerboard (see Methods for more details).

Virtually no unit showed a significant covariation with any of the four predictors at any time during the Targets-observation period of the TF task (Figure 5). In contrast, after checkerboard onset, when the monkeys could determine the checkerboard’s dominant color and choose a reach target, the proportion of units that reflected the direction of the chosen target (green) rose rapidly, and a somewhat smaller proportion of units reflected the signed strength of the color-independent directional evidence towards a given target (blue). In contrast, very few units were significantly modulated by the color of the chosen target (magenta) or the signed checkerboard color coherence at any time after the checkerboard appeared (cyan).

**Figure 5.**
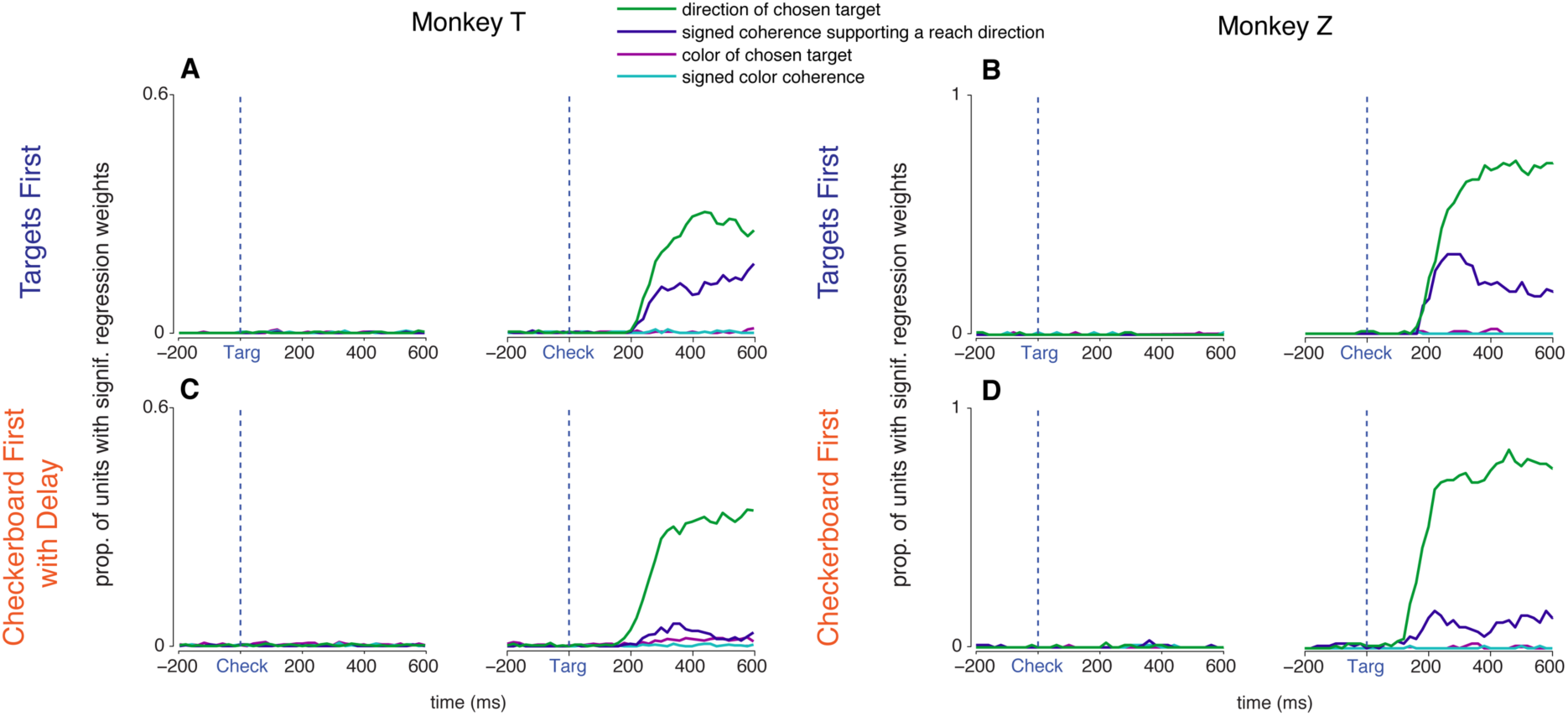
Single-neuron and population-level neural responses reflect chosen reach direction and evidence towards a reach direction, but not chosen target color or signed color coherence of the checkerboards. Each unit’s firing rate at a given 20ms time point across trials was regressed on a linear model with the following predictor variables: direction of the chosen target (green), signed checkerboard coherence favoring a particular reach direction independent of its color (teal), signed checkerboard color coherence (blue), and the color of the chosen target independent of its direction (magenta). This regression analysis was repeated for all units and all time points from −200ms before to +600ms after the appearance of the first visual cue in each trial (left of each pair of figures) and of the second visual cue (right). Plotted are the proportions of units with significant regression weights for a given predictor at each 20ms time point. Very few significant correlations with any regression predictor were seen during the observation period of the first visual cue in either task. Significant correlations with chosen target color (magenta) and signed checkerboard color coherence (blue) rarely occurred at any time in either task in either monkey. However, following the second visual cue, there were significant correlations with the direction of the chosen target (green) and the level of checkerboard evidence favoring a target direction (teal). Correlates with the direction of chosen target were comparably frequent in both tasks for each monkey. However, the incidence of significant correlations with checkerboard evidence favoring a particular reach direction was substantially lower in both monkeys in the CF/CFD tasks than in the TF task.

Similarly, in the CF/CFD task, virtually no unit showed a significant correlation with the color of the chosen target or the signed color coherence of the checkerboards at any time in the trial. After the targets appeared, the proportion of units with effects of chosen reach direction rose rapidly for both monkeys. Significant correlations with signed directional coherence did not appear during the Checkerboard-observation period in either monkey, but did appear after the targets appeared, even though the checkerboard was no longer visible for monkey T. The incidence of directional coherence effects was lower than in the TF task in both monkeys. These single-unit linear regression findings were confirmed and complemented by repeated-measure ANOVA, which found strong main effects of chosen reach direction and unsigned evidence strength after the second cue appeared in each task, and very few main effects of checkerboard dominant color (Supplemental Figure 4; Supplemental Table 1).

In summary, we emphasize three main findings. 1) The chosen direction of movement was by far the strongest predictor of neural activity in both monkeys in both tasks, and was only expressed after the monkeys had received both instructional cues. 2) The next strongest predictor was a color-blind signed signal that instructed the monkeys in which spatial direction to reach; this predictor reflected the combination of information provided by the two visual cues and was graded by the strength of evidence in the checkerboard. This is the component that most strongly implicates PMd in the action-choice process based on sensory evidence. It arose only after the monkeys received both cues, and appeared at the same time as the activity explained by the chosen direction predictor in the TF task. Interestingly, it was also present in the CF/CFD task, but was weaker in both monkeys, suggesting that the evidence provided by the checkerboard was at least partially taken into account by the time the colored targets appeared. 3) There were almost no significant correlations with either chosen target color or signed color coherence of the checkerboards, at any time in either task. That is, there was no prominent neural correlate in PMd of the perceptual deliberation process that was essential to identify the checkerboard’s dominant color and correct target color.

### PMd encoded reach direction sooner in the CF/CFD task than in the TF task

Another important question is whether the order in which the cues were presented had an impact on the onset latency of choice-related activity of PMd neurons in the two tasks. We did an ROC analysis of the ability of an ideal observer to predict the dominant checkerboard color or the chosen target direction based on neural activity in sequential non-overlapping 20ms time bins. Consistent with the regression results, we found strong population-level signals about target choices in both monkeys only after both visual cues appeared in each task (Figure 6). In contrast, the ability of an ideal observer to discriminate the checkerboard’s dominant color remained at baseline during the Checkerboard-observation epoch of the CF/CFD task in both monkeys, and remained at baseline after the second visual cue appeared in both tasks in monkey Z. There was a statistically significant but very modest increase in the discriminability of the checkerboard dominant color in the population activity of monkey T after targets appeared in the CFD task but not after the checkerboard appeared in the TF task (Figure 6).

**Figure 6.**
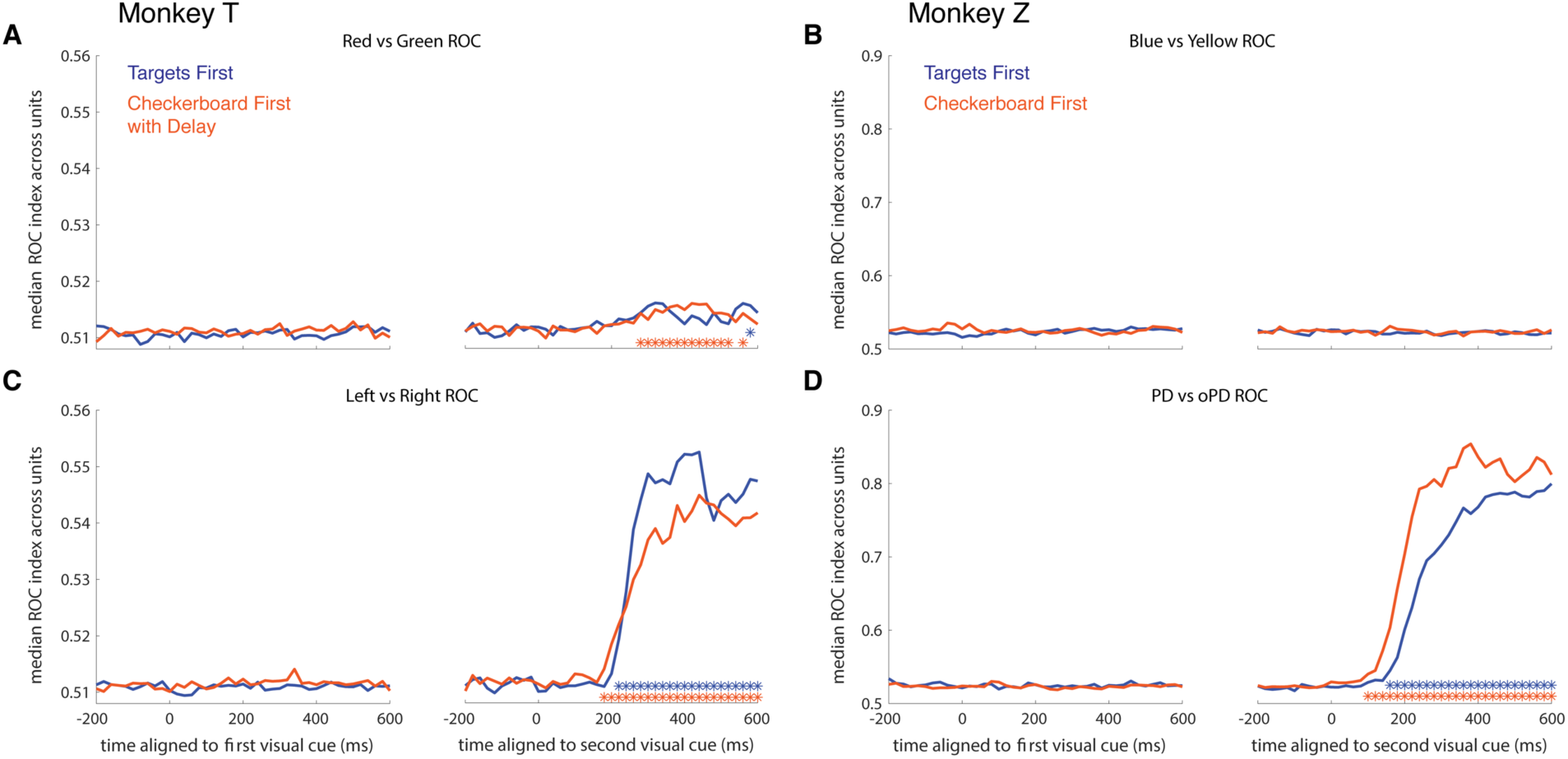
Ideal-observer analysis of the detectability of population-level neural signals about checkerboard dominant color versus chosen direction of reach. Receiver operating characteristic (ROC) analysis was performed on the single-trial discharge rates for each unit at every 20ms time point from −200ms before to +600ms after the appearance of the first (left of each pair of figures) and second (right) visual cue in each trial, for trials sorted according to the dominant color of the checkerboard (**A,B**) or the chosen direction of reach (**C,D**) in the TF (blue) and CFD/CF (orange) tasks. The median area-under-the-ROC-curve (AUC) value across the population of units is plotted at each 20ms time point. The median AUC values for the Color test were very small at all times in both tasks in both monkeys (**A,B**), indicating a very weak representation of information about the dominant color of the checkerboards. In contrast, there was an abrupt increase in median AUC values for Direction shortly after the appearance of the second visual cue in both tasks in both monkeys (**C,D**). To detect a significant increase in the detectability of Color or Direction information in the population activity, the distribution of AUC values measured in each 20ms time step was compared to a baseline 20ms interval −200ms before the appearance of the first visual cue (1-tailed Wilcoxon signed rank test, p < 0.01). Time steps in which the AUC values were significantly higher than the baseline distribution are indicated by blue and red asterisks for the TF and CFD/CF tasks, respectively.

To estimate the onset of a significant increase in the AUC values of the ROC analysis, we used a bootstrap test in 50ms bins after the appearance of the second visual cue in each task compared to a baseline time bin 200ms before the second cue appeared (see Methods). For both animals, the latency for reach direction detection was shorter in the CFD task than the TF task (monkey T, TF task: 208 ± 5 ms, CFD task: 175 ± 14 ms; monkey Z, TF task: 193 ± 9 ms, CFD task: 134 ± 15 ms). These distributions were also highly significantly different (Wilcoxon signed rank test, monkey T: p = 2.9e-165, monkey Z: 2.11e-167).

These shorter onset latencies in the CF/CFD task than the TF task further support the conclusion that some information about the checkerboard color composition had been processed during the Checkerboard-observation period in both monkeys, and suggests that the temporal dynamics of neural responses in PMd following the appearance of the second visual cue are different in the two tasks, resulting in a reduction of the onset latency of the earliest PMd activity predicting action choices in the CF/CFD task.

## Discussion

Our goal was to investigate how the demands of a task affect the extent to which neural correlates of perceptual decision-making are present in dorsal premotor cortex (PMd). In many studies of decision-making, the action options to report a given decision are specified before the decision cue is presented (Kim and Shadlen, 1999; Roitman and Shadlen, 2002; Shadlen and Newsome, 2001). Similarly, in this study, the appearance of the two color-coded target cues at the start of each trial in the TF task provided the specific evidence-report mapping before the colored checkerboard cue appeared. In contrast, in the CF/CFD task, we inverted the task timeline by presenting the colored checkerboard before presenting the two color-coded targets. Each cue provided different partial information about the required reach movement, and the monkeys could not choose a specific reach action until it received both pieces of information.

We found that, consistent with previous results (Chandrasekaran et al., 2017; Coallier et al., 2015), PMd activity reflected decision-related correlates after checkerboard onset in the TF task, including the level of the color-independent checkerboard coherence supporting a target in a specific location (signed directional coherence) and the ultimate action choice, but not the physical color composition (signed color coherence) of the checkerboard or the color of the chosen target. Similar response effects were observed in the CF/CFD task after the colored targets appeared. Importantly, no differential decision-related activity that predicted any aspect of the monkeys’ perceptual or action-related choices was expressed in PMd while the monkeys observed the first cue in each task. The latter findings are particularly striking, since many PMd neurons in monkey Z emitted overt but non-differential responses to the first cue in each task, and monkey T had to retain some form of information about the color composition of the checkerboard in memory before the colored targets appeared in the CFD task.

### PMd’s role in perceptual decision-making

Perceptual decision-making has been described as the process by which an individual commits to a proposition on the basis of perceived sensory information; that decision is typically reported in animal studies by a differential motor response (Gold and Shadlen, 2007; Shadlen and Kiani, 2013). Of course, as already noted, we make many perceptual decisions every day (“I like those boots”) without committing to a particular action, unless one counts that inaction as a motor decision. In this study, a categorical perceptual decision about the dominant color of a checkerboard stimulus is combined with the information about the spatial configuration of colored targets to arrive at a motor decision. In the TF task, the nominally perceptual decision about the checkerboard is made in the cognitive context of a priori known evidence-report mappings provided by the colored target cues. In contrast, the CF/CFD tasks were designed to permit a non-motoric perceptual decision about the checkerboard before the associated evidence-report mapping required to make a motor decision is known. The critical novel finding of the present study is that the appearance of the checkerboards in the CF/CFD task did not elicit any activity in PMd in monkey T and no color-specific correlates of either the sensory evidence (signed color coherence) or a categorical decision about the dominant color during an ongoing perceptual decision process in monkey Z. These results confirm that PMd is implicated in the processing of sensory information supporting reach action choices and reveal that PMd primarily processes the spatial information provided by the instructional cues about action choices. Furthermore, it expresses correlates of the differential action-related decision process only after the monkeys have received all the information from both cues required to map checkerboard color information onto a specific target choice. The near absence of significant correlates of the critical physical property of the cues – color - that informs the final action choice indicates that it does not make a substantial contribution to the non-motor perceptual aspects of these tasks. These findings also constrain how PMd is implicated in the motor decision. The PMd neurons may be receiving a time-varying signal about the most likely color of the checkerboard but are directly transforming that information into a signal about the most likely spatial location of the target, once their color-location conjunctions are known. Alternatively, the PMd neurons may be receiving a more abstract signal about the mounting evidence for the solution of the color/location matching rule based on the color information in the checkerboard, and are translating that into a signal about the most likely target location for a reach independent of its color. Consistent with this hypothesis, Mante et al (2013) have shown that prefrontal cortex neurons in and near the frontal eye fields encode the color information in colored RDK stimuli and convert it into evidence for the choice of a color-coded saccade target in a task that is conceptually similar to the TF task. Similarly, we have preliminary evidence that dorsolateral prefrontal neurons near the principal sulcus signal the color/location conjunctions of the targets during the target-observation period and the mapping of the checkerboard dominant color onto the color-matched reach target after the checkerboard appears in a TF task (Coallier et al., SfN abstract 2008). Finally, PMd may be receiving a time-varying signal about the most likely location of the target after all salient information has been processed and is only implicated in the preparation of the chosen action. However, this last possibility seems to be unlikely given all the previous studies that have implicated PMd in the conversion of sensory information into abstract goals and specific actions (Boussaoud and Wise, 1993; Chandrasekaran et al., 2017; Cisek and Kalaska, 2002, 2005, 2010; Coallier et al., 2015; Crammond and Kalaska, 2000; di Pellegrino and Wise, 1991; Hoshi and Tanji, 2000, 2006; Kalaska and Crammond, 1995; Nakayama et al., 2008; Thura and Cisek 2014; Wallis and Miller, 2003; Wise et al., 1992, 1997).

### Comparison of findings in the two monkeys in the TF, CF and CFD tasks

The behavioral and neurophysiological results were very similar in both monkeys in the TF task. However, monkey T’s behavior in the CFD task was inconsistent with that of monkey Z and human subjects in a CF task (Coallier and Kalaska, 2014). Its RTs were longer than in the TF task, especially for the checkerboards with the strongest color coherences, and monkey T began to make target-choice “lapse” errors to those same stimuli. This was accompanied by slower rates of growth of the choice selectivity signals in PMd for higher-coherence checkerboards in the CF task than the TF task (Figure 4). There are several possible contributing factors: 1) the 500ms checkerboard observation period may have been too short to complete a categorical perceptual decision on every trial while it was still visible; 2) the checkerboard observation period may have been too short to form an accurate short-term memory of its physical features during the 400-800ms delay period; or 3) the memory of the checkerboard or of the perceptual decision may have decayed during the delay period. Monkey T’s success rates in the CFD task did improve by ∼5% for shortest compared to longest delay periods, as well as when longer checkerboard durations were tested (data not shown).

Furthermore, monkey T may have adopted a strategy of trying to store a memory trace of some features of the checkerboard and largely deferring the decision process until the colored targets appeared in the CFD task. A perceptual decision might be formed in CFD-like tasks by sequential sampling from working memory (Shadlen and Shohamy, 2016); some preliminary evidence supports this hypothesis (Shushruth and Shadlen, CoSyNe abstract). Alternatively, monkey T may have at least partly formed a perceptual decision before the targets appeared, but the dynamics of its motor decision were still sensitive to the strength of the sensory evidence that had informed the perceptual decision, for instance by being modulated by monkey T’s level of confidence in the correctness of its perceptual decision (Drugowitsch et al., 2014; Kiani and Shadlen, 2009; Kiani et al 2014; Zylberberg et al., 2016).

In contrast, Monkey Z showed nearly identical psychophysical performance in the TF and CF tasks and the same chronometric savings in the CF task as the human subjects (Coallier and Kalaska, 2014), suggesting that it had largely or completely processed the color evidence in the checkerboards before the targets appeared. This was accompanied by higher rates of rise of choice selectivity signals for the 4% checkerboard coherences in the CF task than in the TF task (Figure 4). Monkey Z continued to show a large reduction of RTs for checkerboards with low color coherences in a CFD task (Supplemental Figure 1). Nevertheless, it did show some of the same costs as monkey T for the brief fixed checkerboard observation period and memory delay; its RTs for the strongest checkerboard color coherences were modestly prolonged by ∼25ms compared to that in the CF task, and it had a lapse-error rate of ∼10% for high-coherence stimuli. These costs in both monkeys are somewhat surprising. Easy color discriminations such as those for checkerboards with 80-100% coherence should be very rapid (30-50ms; Stanford et al., 2010; Shankar et al., 2011). Similarly, the time needed for PMd neurons to express differential action choices based on the color of an instructional cue (e.g., go/nogo; reach toward or away from a visual cue) likewise takes only ∼50-100ms (Crammond and Kalaska, 1994; Kalaska, 1996; Kalaska and Crammond, 1995). The imposed delay period seems to have impeded the ability of both monkeys to retain information about the dominant color of the checkerboard or to use it correctly to identify the reach target after they appeared. This effect was most apparent for the nominally “easiest” checkerboards in both monkeys.

Many PMd neurons in monkey Z also responded to the appearance of the first cue in each task (Figure 2, 3, Supplemental Figure 3). These responses may contribute to the overall target selection process (Cisek and Kalaska, 2005, 2010), but they did not encode the color information in either the targets (TF task) or the checkerboard (CF task), or display any differential decision-making signals that predicted either the target color or direction choices of the monkey (Figure 4-6). Thus, unlike monkey T, monkey Z’s performance of this task involved engagement of PMd even when only partial information was available about the specific action choices, but which may reflect the likelihood of potential reach directions. This is consistent with prior reports of non-differential activation of premotor cortex neurons before the final action was specified (Cisek and Kalaska, 2005; Hoshi and Tanji, 2006; Nakayama et al., 2008; Yamagata et al., 2009, 2012). For instance, PMd activity can signal the likely next direction of movement as a function of current arm position within a limited task workspace before the next target appeared in a random-walk reach sequence task (Glaser et al., 2018).

Despite these differences in behavior and neural responses between the two monkeys, our primary findings that PMd neurons did not strongly encode the critical color dimension of the instructional cues and did not show neural correlates of differential decision-making processes until the monkeys had received the information provided by both instructional cues were remarkably robust in both animals. Furthermore, both monkeys showed a reduction in the prominence of neural correlates with signed directional coherence evidence in the CF/CFD task compared to the TF task and an earlier appearance of choice-related activity in the CF/CFD task than the TF task. This suggests a difference in the temporal dynamics of decision-making in the two tasks and at least some degree of processing of the checkerboard sensory evidence in the CFD task before the targets appeared in monkey T, despite the lengthening in RTs.

## Comparison with other decision-related areas

Other motor areas such as the superior colliculus (SC; Horwitz et al., 2004), presupplementary and cingulate motor areas (Merten and Nieder, 2012), and especially lateral intraparietal cortex (LIP) (Bennur and Gold, 2011; Freedman and Assad, 2006) have shown different degrees of correlations to perceptual decisions versus motor actions, by presenting the sensory evidence before the target choices, and by decoupling the mapping between the evidence and action choices. For instance, by inducing saccades with intracortical microstimulation, Gold and Shadlen (2003) found behavioral evidence of developing oculomotor commands in the frontal eye fields (FEF) directed towards known target locations either in the direction of RDK motion (pro-saccade task) or the opposite direction (anti-saccade task) while the monkeys observed RDK stimuli, but not in a colored-target task in which red and green targets appeared at random locations only after the RDK stimuli were extinguished. The results implicated FEF primarily in the motor aspects of the sensorimotor decision process, and only when the evidence-action associations were known, consistent with the present findings. Horwitz et al (2004) presented report saccade targets at unpredictable locations after RDK stimuli were extinguished. Some SC saccade-related neurons began to discharge during RDK motion; their activity reliably predicted that the monkey would ultimately choose the target that signified perceived RDK motion towards the neuron’s preferred movement field, even though the report saccade direction was not yet known and could be in a very different direction. They suggested that this activity may have been part of a mnemonic representation of the sensory evidence on which the monkeys would base their motor decision, expressed in the spatial framework of saccade movements in SC circuits. Finally, Bennur and Gold (2010) used a variant of the colored-target task (Gold and Shadlen, 2003) in which they presented the spatial locations of red and green targets before (Task 1), during (Task 2) or after (Task 3) the RDK stimuli were presented. Tasks 1 and 3 are conceptually similar to our TF and CF tasks, respectively. They found that many LIP neurons showed significant correlations with different combinations of RDK motion direction and strength, target color, and saccade direction choice at different times in each trial. Critically, they found that saccade-related LIP neurons robustly encoded the target colors and the perceived RDK motion direction after each stimulus was presented even if the information provided by the other stimulus was not yet available, and strongly represented the chosen saccade direction after both stimuli had been presented. Thus, LIP neurons could express neural correlates of all salient sensory and motor aspects of the sensorimotor decision leading to saccade direction choices, including a representation of perceived RDK motion direction before the metrics of the report saccades were known (c.f., Fanini and Assad, 2009). These findings of Bennur and Gold (2011) are in stark contrast to our own. Finally, Freedman and colleagues likewise found explicit representations of sensory properties, cognitive decisions and motor reports in LIP (Freedman and Assad, 2006, 2010; Ibos and Freedman, 2017). These results suggest that the parietal cortex may be tightly involved at the intersection of sensory and motor processing, even representing the salient sensory evidence explicitly in its activity, whereas PMd does not in our tasks.

### Comparison to previous PMd findings

Our findings are consistent with previous studies showing that PMd activity reflects sensory cues that guide motor goals. Wise and colleagues (Boussaoud and Wise, 1993; Wise et al., 1992) showed that the responses of PMd neurons to a visual instructional stimulus are strongly modulated when it signals different motor responses or a shift in attention rather than a movement. Di Pellegrino and Wise (1991) recorded from PMd in an instructed-delay match-to-sample task similar to the CFD task in which the color (R/G) of the first cue signaled which button to press after colored stimuli appeared above them. They found that ∼11% of PMd neurons responded to the initial cue, similar to monkey Z, but did not signal the color of either the first cue or the selected button, whereas many neurons signaled which button was pressed, similar to both monkey T and Z. In contrast, some dorsolateral prefrontal cortex neurons showed correlates with the colors of the cues in the same task. Many PMd neurons can signal whether a future reach will be to the leftward or rightward of two targets independent of the physical identity of the instructional cues, even if the subject does not know exactly where on the screen the targets will appear (Nakayama et al., 2008). One possibility is that PMd represented this abstract goal via spatial encoding, similar to a spatial mnemonic strategy suggested by Horwitz et al. (2004). This may also be a more abstract form of the representation of the spatial location of potential reaching targets in a 2-Target instructed-delay task (Cisek and Kalaska, 2005; Coallier et al., 2015; monkey Z in the present study). Neural responses in the 2-Target task also showed only very modest correlations with the colors of the instructional cues (Cisek and Kalaska, 2005; Coallier et al., 2015). All of these findings indicate that PMd is predominantly implicated in processing the spatial information about future action choices provided by instructional cues, but does not strongly represent the salient physical properties of the stimuli. Wallis and Miller (2003) trained monkeys to report whether two sequential complex visual images that changed each day were similar or different by releasing a key with their hand either immediately after receiving the second image, or after a further 500ms delay. A “rule” cue presented at the same time as the first visual image instructed the monkey to release the key immediately if the two stimuli were either the same or different, in each trial. Very few PMd neurons differentially encoded the identity of the visual images. However, 48% of the PMd neurons differentially signaled the same/different rule in each trial that defined the stimulus conditions required to release the key immediately or after a delay, and 72% signaled the motor decision to release the key immediately or later after the second image appeared. Thus, unlike our CF/CFD task, their monkeys always knew the report action to take (release the key) and the sensory evidence only determined when to execute that action.

Finally, Romo and colleagues have studied PMd in tasks in which monkeys must push the leftward or rightward of two closely-spaced buttons with one hand to report whether the first or second of two sequential tactile stimuli delivered to the hand of the other arm was of a higher relative vibration frequency (Romo et al., 2004; Romo and de Lafuente, 2013) or whether they had the same or different temporal structure (Rossi-Pool et al., 2016; 2017). In the vibratory-frequency task, few PMd neurons (∼10%) showed correlates with the frequency of the two stimuli, and even fewer correlated with the chosen action, suggesting that the sample population was not strongly implicated in either the perceptual or motor aspects of that task. In contrast, ∼20-35% of a different PMd sample population in the second study expressed a differential categorical signal about whether the first tactile stimulus had a pattern of constant inter-pulse intervals or contained a high-frequency burst when it was presented, others signaled the specific ordinal sequence of the different stimulus pairings after the second stimulus was presented, and ∼20-30% signaled the categorical decision of whether the two stimuli had the same or different temporal structure independent of their actual structure. The last neurons could also be a potential correlate of the final motor decision because of the fixed relationship between same/different and left/right buttons, reminiscent of the findings by Nakayama et al. (2008). The Rossi-Pool et al. (2016, 2017) study reported a much stronger representation of the salient physical attributes of the sensory cues in PMd than in the CF/CFD tasks in our study, but unlike the present study, the sensory cues in the temporal-structure comparison task were received in a context in which the ultimate report actions were fixed and known in advance.

### Summary

When the perceptual assessment of a checkerboard decision cue could be made in the context of known specific evidence-report mappings onto action choices (TF task), dorsal premotor cortex neurons generated a differential decision-making signal reflecting the strength of the sensory evidence supporting the spatial location of the target and the final action choice, but not the critical physical dimension of the checkerboard, its dominant color. When this link was broken and perceptual decisions about the checkerboard could be formed before the specific action to report the decision was known (CF/CFD task), PMd did not respond in a differential decision-related manner to the checkerboard. Explicit representations of the color/location conjunctions of the targets or of the color composition of the checkerboard, that are required to perform the tasks, were not found in PMd, and are presumably expressed elsewhere. Dorsolateral prefrontal cortex is a leading candidate (Di Pellegrino and Wise, 1991; Mante et al., 2013), and is currently under study in these tasks.

## Acknowledgments

Support for this project was provided by the following sources:

MW: US DoD, Air Force Office of Scientific Research, National Defense Science and Engineering Graduate (NDSEG) Fellowship, 32 CFR 168a. CM, JFK: Canadian Institutes of Health Research operating grants MOP-97944 and MOP-142220, and an infrastructure grant from the FRSQ to the Groupe de recherche sur le système nerveux central. CC: NIH/NINDS K99/R00 grant NS092972 and Howard Hughes Medical Institute. DP: the Champalimaud Foundation, Portugal, and Howard Hughes Medical Institute. KVS: NIH/NINDS Transformative Research Award R01NS076460, NIH/NIMH Transformative Research Award R01MH09964703, NIH Director’s Pioneer Award 8DP1HD075623, DARPA Biological Technology Office (BTO) “REPAIR” award N66001-10-C-2010, DARPA BTO “NeuroFAST” award W911NF-14-2-0013, the Simons Foundation Collaboration on the Global Brain awards 325380 and 543045, and the Howard Hughes Medical Institute. We thank Dr. William T. Newsome for comments on the manuscript and discussions throughout the course of this work.

## Author Contributions

Study conception and design: MW, CM, CC, KVS, JFK.

Data acquisition and analysis: MW, CM, CC, JFK.

Manuscript preparation and editing: MW, CM, CC, DP, KVS, JFK.

Supervision of project: KVS, JFK.

## Supplemental Tables

## Methods

### General Information

Two rhesus macaque monkeys (*Macaca mulatta*), monkey T (9-year-old male, 15.0 kg; same monkey T as in Chandrasekaran et al., 2017) and monkey Z (9-year-old male, 12.0 kg), were used in the study. Monkey T’s home environment, standards of care, and this experiment were approved by the Stanford University Institutional Animal Care and Use Committee. Monkey Z’s housing, veterinary care and experimental protocols were approved by the institutional animals-in-research committee (CDEA - Comité de déontologie de l’expérimentation sur les animaux, *Université de Montréal*), and respected all institutional and national guidelines.

### General task design

We used two main variants of a decision-making task in which subjects chose between two opposite reach directions based on the content of two successive visual stimuli that provided different types and amounts of sensory evidence supporting each reach choice in each trial. In both variants, the goal for the subject was to determine the dominant salient color of a checkerboard-like visual stimulus, and to report that color by making an arm reach to the corresponding colored target. The difficulty of the sensorimotor decision was manipulated by varying the relative numbers of squares of two task-relevant colors. This was roughly equivalent to varying the coherence of dot motion in RDK tasks. However, RDK tasks involve detecting a coherent-motion signal against a random-motion noise background. In contrast, in the checkerboard stimuli, each evidence element (a colored square) is easily detected and discriminated. The challenge is to assess their relative numbers to estimate the dominant color of the checkerboard. We use the term “color coherence” here to indicate the degree to which the squares in the checkerboard are the same color or not. Thus, if all the squares are of the same color, the color composition of the checkerboard stimulus is said to be 100% coherent whereas a checkerboard with equal numbers of the colored squares has 0% color coherence.

A key differentiator for this study compared to RDK tasks is that the color of an object does not have any inherent association with any parameter of a reach movement, such as target spatial location or reach direction. Color only becomes action-relevant in our tasks by application of an arbitrary stimulus-response mapping rule; the subjects decide on the dominant color of the checkerboard and then use an operantly-conditioned color-location matching rule to associate it with the target of the same color. This is not the case in RDK stimuli, which have an intrinsic physical property — the spatial direction of coherent dot motion — that is also usually directly mapped onto the direction of the motor output. A second differentiator is that typical RDK stimuli are stochastic and dynamic, with variable numbers of short life-time dots moving in the coherent and random directions from frame to frame. In contrast, all checkerboard stimuli observed by monkey T and half the stimuli presented to monkey Z were static, with each square remaining visible and stationary for the entire duration of the checkerboard observation period. A third differentiator is that low-coherence RDK stimuli contain very few dots that move coherently in the task-salient direction and the primary perceptual challenge is detecting that weak signal in the random-motion noise. In contrast, low-coherence checkerboards contained large but nearly equal numbers of easily discriminable colored squares that each unequivocally supported one or the other of the two action choices.

The Targets First (TF) task variant followed the event timeline used in many sensorimotor decision tasks (Chandrasekaran et al., 2017; Coallier and Kalaska 2014; Coallier et al., 2015; Kim and Shadlen, 1999; Shadlen and Newsome, 2001). First, two color-coded targets appeared, providing the subject with sensory information about the two reach choices (Cisek and Kalaska, 2005) and how color (red or green for monkey T, blue or yellow for monkey Z) would be associated with reach direction. The checkerboard appeared later. Deliberation about dominant checkerboard color could occur concomitantly with planning for the reach, because each color was already associated with a specific target location. The monkeys were free to initiate a reach to a target at the time of their choosing after checkerboard appearance.

Crucially, in the Checkerboard First (CF) and Checkerboard First with Delay (CFD) tasks, the order of the two sensory events was reversed. The checkerboard appeared first, but the monkeys did not yet know which color would be associated with a given target location and reach direction. Thus, the monkeys could in theory deliberate upon the checkerboard’s dominant color, but could not prepare a specific motor response to report it.

Details of task structure and recordings varied between the two laboratories, as detailed below.

### Monkey T method details (Stanford University) Experimental setup

Throughout the experiment, monkey T sat in a primate chair (Crist Instruments, Snyder Chair) ∼30 cm in front of an LCD computer monitor (Acer HN274H, then Acer ×G270HU) on which the task would be presented. The animal’s non-reaching (left) arm was loosely restrained with a tube and cloth sling. The stimulus presentation and data collection were controlled by a custom computer system (MathWorks’ xPC Target and Psychophysics Toolbox). We placed a photodetector (ThorLabs PD360A) in the corner of the computer screen to detect the onset of various task events to a 1 ms resolution. Hand position was measured by taping a reflective bead (11 mm, Northern Digital Inc’s Passive Spheres) to the tip of the middle finger of the reaching (right) hand, and tracking the location of this bead in three-dimensional space using an infrared tracking system (Polaris Spectra; Northern Digital Inc.). Eye position was tracked using an infrared camera (ISCAN ETL-200 Primate Eye Tracking Laboratory) mounted overhead; the eye image was reflected to the camera above using an infrared mirror (ThorLabs) placed at a 45° angle in front of the animal’s nose. The infrared mirror allows visible light to pass through, so it does not obstruct the animal’s view of the computer monitor.

### Task design

In the Targets-First (TF) task (Figure 1A), monkey T initiated a trial by placing its right hand on a center hold circle (24 mm diameter) and fixating its gaze on a cross (6 mm diameter), located above the center hold circle. Once these two conditions were met, there was a brief variable delay of 250-400 ms, and then two monochromatic targets (one red, the other green) were presented 100 mm to the left and right of the center hold. These targets were presented for 450-800 ms, during which the animal maintained center hold and eye fixation (“Targets-observation period”). Finally, a static checkerboard stimulus containing variable numbers of red and green squares from trial to trial was presented, centered at the fixation cross, and served as the go cue for monkey T to make its report (“Checkerboard-RT epoch”). It was free to initiate its chosen reach action as soon as it was ready. The moment that center hold or eye fixation was broken at the onset of its reach response, the checkerboard disappeared but the targets remained visible.

In the Checkerboard-First with Delay (CFD) task (Figure 1A), the presentation order of the targets and checkerboard was reversed. The trial began in the same way as in the TF task, and the center hold delay was the same at 250-400 ms. Then, the checkerboard appeared for a fixed period of 500 ms (“Checkerboard-observation period”), and subsequently disappeared for a memorized-delay period of 400-800 ms, after which the colored targets appeared. The appearance of the targets served as the go cue for Monkey T to make its report (“Targets-RT epoch”). As in the TF task, the monkey could initiate its reach movement as soon as it made its target choice.

The size and locations of all visual stimuli were the same in both tasks, and the red and green colors for the targets and checkerboard were identical and isoluminant (22 cd m-2, Konika Minolta). The assignment of red and green to left and right targets was randomized between trials. The checkerboard consisted of a 15 x 15 grid of 2.5 mm x 2.5 mm squares. The task difficulty was adjusted by varying the number of red and green squares in the checkerboard (Figure 1D). For each dominant color (red or green), we used seven difficulty levels (# non-dominant squares and # dominant squares: 11+214, 45+180, 67+158, 78+147, 90+135, 101+124, and 108+117). These levels correspond to coherence levels (difference in red and green squares, divided by the total number of squares) of 90.2%, 60%, 40.4%, 30.7%, 20%, 10.2% and 4%. In each trial, a single static checkerboard matrix was presented in which the R and G squares were distributed randomly within the 225-square checkerboard matrix. A different random matrix was presented on each trial, even within the same checkerboard coherence. All task factors (dominant checkerboard color, checkerboard coherence, and correct colored target location) were presented in a randomized sequence. Data were collected until neural isolation was lost or until the monkey was sated. Typical daily data sets comprised roughly 2000 trials of CFD only, or 1000 trials each of CFD and TF.

### Training history

Monkey T was first trained to make reaching movements towards targets on the computer screen, for pieces of fruit and then for juice reward. It was then trained on the TF task, starting with the highest checkerboard coherence (90.2%). At the beginning, a high-coherence checkerboard was presented before the targets were presented; once the association between checkerboard color and target color was learned, the order was inverted to present the targets first. Further details can be found in (Chandrasekaran et al., 2017). To train monkey T on the CFD task, we began by presenting only the highest checkerboard coherence with a 300 ms delay between checkerboard presentation and target presentation. We gradually increased the delay and added gradually lower checkerboard coherences, over many daily training sessions. Monkey T did not experience targets at any location other than left or right of the central start position.

### Recording chamber implantation and neural data collection

Monkey T had an acrylic head implant with a recording chamber over left PMd/M1 (coordinates A16, L15; Figure 1E)). In this recording chamber (19 mm diameter), a series of small burr holes (3 mm diameter) were drilled through the acrylic implant and skull to access dura and brain. The neural data were recorded using either single electrodes (22 sessions) or linear arrays (19 sessions). Single electrodes were FHC tungsten electrodes #UEWLGCSEEN1E (Frederick Haer & Co, Bowdoin, ME USA). Linear arrays were Plexon (Dallas, T× USA) U probes with platinum-iridium recording sites, 16 channels spaced 150 um apart (specifically: PL×-UP-16-15ED-150-SE-100-25(640)-15T-700). Single electrodes were lowered into the brain until a unit was found; linear arrays were lowered until all electrode sites were in brain, preferably with a unit on the deepest and shallowest electrodes. The units were sorted online using BlackRock Central software. Units were included in analysis if they were responsive at any time during the trial. Data were collected in blocks of roughly 500 trials per task.

### Neural data pre-processing

To identify putative single units, we examined the inter-spike interval (ISI) distributions for each unit (Hill et al., 2011). Spike timing information was collected at 30,000 samples / s. We considered ISI violations to be those ISI that were less than 2 ms, a conservative refractory period between action potentials. A unit was considered a single unit if it had less than 1.5% ISI violations. Of the 499 units collected in CFD task, 441 (88.4%) units were identified as single neurons, with a mean of 0.44% ISI violations; the remaining multi-units had a mean of 3.48% ISI violations. Of the 351 units collected in TF task (all of which overlap with the CFD units), 310 (88.3%) units were identified as single units, with a mean of 0.46% ISI violations; the remaining multi-units had a mean of 2.72% ISI violations. Of these units, 304 were consistently classified as single unit across both tasks and 33 were consistently classified as multi-unit across both tasks.

Firing rates were constructed by convolving a 50 ms acausal box car filter with spike times at 1 ms resolution. The exception is for Figure 2, in which we used a 75 ms box car filter for better visualization. Behavioral reaction time in each trial was calculated as the time at which hand velocity exceeded 10% of the maximum hand velocity. Data shown include correct and incorrect trials (depending on the analysis), with reaction times greater than 300 ms, and do not include overt “change-of-mind” trials in which the monkey began to reach to one target and then reversed direction and completed a reach to the opposite direction (Resulaj et al 2009; Kaufman et al 2015).

### Monkey Z method details (Université de Montréal) Experimental setup

Where monkey T used its arm to touch targets on a monitor screen, monkey Z used a pendulum-like handle that moved over a horizontal digitizing tablet (hand position measurements at 100 Hz, ±0.05 mm precision; for technical details of the task apparatus, see Coallier and Kalaska, 2014), to displace a 6 mm cross-shaped cursor between targets displayed on a vertical computer monitor at a viewing distance of 60cm.

### Task design

The TF task structure for monkey Z was identical to the “Choose-and-Go” task used in previous studies (Coallier and Kalaska, 2014; Coallier et al., 2015). Each trial began when a small open white square (1.0 cm) appeared at the center of the monitor (Figure 1A). The monkey used its arm and pendulum to position the on-screen cursor in the central square and held it there for 500 ms. Two monochromatic square target cues (4.5 cm; one yellow and one blue) then appeared at opposite sides of the central square (15.5 cm separation between the centers of the target squares) for 1250±250ms (Targets-observation period). The same two opposite target locations (out of 8 possible locations arranged in a circle) were used for each block of trials for a given neuron according to its reach-related directional tuning, but varied from neuron to neuron (see below). After a variable period of 1250±250ms, the central square was replaced by the checkerboard stimulus, and white squares appeared at the other 6 target locations in the task, serving as the go signal (Checkerboard-RT epoch). Monkey Z was free to reach to the chosen target at any time, without an imposed pre-reach delay. The checkerboard stimulus disappeared as soon as the cursor position exited the boundary of the original small central target. Monkey Z had to reach the target within 750ms and stay within the target for 1000ms to receive a liquid reward if the chosen target was correct.

The checkerboard consisted of a 15 x 15 grid (4.0 cm) that contained a total of 100 yellow and blue squares plus 125 task-irrelevant red background squares (Figure 1D). For each dominant color (B or Y), three difficulty levels were used (0+100, 40+60, and 48+52) during neural recordings, corresponding to 100%, 20%, and 4% levels of checkerboard coherence. In half of the trials, checkerboards were static, while in the other half they were dynamic – a new checkerboard matrix with the same numbers of colored squares but different square positions was displayed every 50ms. Static versus dynamic stimuli had little or no systematic impact on the psychophysical performance of human subjects (Coallier and Kalaska, 2014) or on the task performance and neural activity recorded in two other monkeys (Coallier et al., 2015). All task factors (correct target location, correct target color, checkerboard color coherence, and static/dynamic checkerboards) were presented in a fully balanced randomized-block sequence. A full task file comprised 120 correctly-performed trials (2 targets x 2 colors x 3 coherence levels x 2 checkerboard conditions x 5 replications). If the monkey chose the incorrect target in a given trial, that trial was re-inserted into the remaining pseudo-random trial sequence until all combinations of trial conditions were completed successfully, resulting in data files containing 120 correct trials and variable numbers of incorrect trials.

Monkey Z performed a version of the Checkerboard First task without a memorized-delay period (CF task, Figure 1A). Its temporal structure was identical to the TF task here, except that the checkerboard cue was presented first during a Checkerboard-observation period after the initial center start-target period. At the end of that observation period, the two color-coded target cues appeared on opposite sides of the checkerboard cue, along with white target squares at the 6 other target locations (Targets-RT epoch). The monkey could initiate its reach choice as soon as it had made its decision. The checkerboard disappeared as soon as the cursor exited the boundary of the original small central target. Data file structure was identical to the TF task.

No gaze fixation control was imposed at any time in any of the tasks for monkey Z (Cisek & Kalaska 2002, 2005; Coallier et al. 2015).

Monkey Z also performed a memorized instructed-delay task with a single target cue presented at the beginning of the delay period (1-Target task, 1T; (Cisek and Kalaska, 2005; Coallier et al., 2015), and a memorized instructed-delay task in which two color-coded potential target cues were presented simultaneously in opposite directions in each trial, followed by a monochromatic central color cue that unambiguously signaled the correct target in each trial (2-Target task, 2T; (Cisek and Kalaska, 2005; Coallier et al., 2015). Monkey Z also performed a version of the TF task that included an extra imposed pre-reach delay period. In this Targets-first with Checkerboard-Delay (TFCD) task, each trial began like the TF task. However, at the end of the initial target observation period, the Checkerboard cue appeared for 1750 ms±300ms, while the target cues remained visible. The monkey was not allowed to make a reach during this Checkerboard-observation period. At the end of that pre-reach delay period, white squares appeared at the other 6 target positions as a go signal and the monkey could make its chosen reach movement. Data file structure was identical to the TF task. Neural data collected from these three tasks will not be presented here.

### Training history

Monkey Z was first acclimatized to sit in a custom-made primate chair, and then trained in a standard 8-direction center-out reaching task without delay periods, for juice rewards. It was then trained to perform the 1T task, followed by the 2T task described before. Following this training, we sequentially introduced the TF, CF and TFCD task variants using multi-colored checkerboard stimuli (Figure 1D). In each task variant, monkey Z first performed the tasks with only the 100% checkerboards, followed by the 20% and then the 4% checkerboards as performance improved and stabilized. After first learning the tasks with the right arm, neural data were collected from the left PMd/M1. Monkey Z was then trained to perform the tasks with the left arm and neural data were collected from the right PMd/M1.

To facilitate comparison of task performance of the two monkeys, monkey Z was also tested in the TF and CF tasks with 7 checkerboard coherence levels (4%, 10%, 20%, 30%, 40%, 60% and 80%) in daily sessions separate from neural recording days. When performing these extended TF and CF tasks (Figure 1B,C), monkey Z performed ∼400 correct trials/checkerboard coherence plus variable numbers of error trials in each task over the course of several daily testing sessions, resulting in ∼6000 trials per task.

To test whether the difference in task performance between the two monkeys in the CF versus CFD tasks was due to the difference in their temporal structure, monkey Z was re-trained and tested in a modified CF task with identical temporal structure to the CFD task, including a fixed 500ms checkerboard observation epoch followed by a variable 400-800ms memory-delay period. Trials were presented with 7 checkerboard coherence levels (4%, 10%, 20%, 30%, 40%, 60% and 80%; Supplemental Figure 1). These behavioral data were collected after all neural data had been collected in the TF and CF tasks.

### Recording chamber implantation and neural data collection

Prior to surgical preparation for neural recordings, an anatomical MRI scan was made of monkey Z’s head to provide images of the sulcal patterns of its cerebral cortex and their location relative to small fiducial-marker gold pins (Hybex Innovations) implanted in its skull at known stereotaxic coordinates. Monkey Z then had custom-made titanium recording chambers implanted over 18 mm diameter trephine holes made in the skull under stereotaxic control (coordinates A21, L15; Figure 1E). The initial implant was over left PMd/M1. After neural data collection was completed, that chamber was removed and the skin opening was closed for several months while it was trained with the left arm. After training with the second (left) arm, a second chamber was implanted over the right PMd/M1. All surgical procedures were performed using standard aseptic surgical procedures (Kalaska et al. 1989; Sergio et al. 2005).

Neural activity was recorded using single in-house-made Corning glass-insulated platinum-iridium microelectrodes. In each daily recording session, the electrode was lowered through the dura and into the brain at a chosen electrode location within the chamber, using a Chubbuck electromechanical microdrive (Georgopoulos et al., 1982). The electrode was advanced while monkey Z performed the 1T and 2T tasks using 8 reach target directions, to search for neurons that showed strong and directionally-tuned activity in the tasks. Once a task-related neuron was isolated, the target that elicited the strongest task-related activity changes was designated as its preferred movement direction (PD). Data were then collected for short trial blocks (20-40 trials) in the PD and the opposite target direction (oPD) in the 1T and the 2T tasks. Data were then collected in the TF, CF and TFCD tasks in pseudo-random order, to collect at least one and ideally two complete data files in each task. A neuron was retained for detailed analysis if its isolation and task-related activity remained stable throughout the recording session, and data were collected successfully from at least one complete data file for the TF and CF tasks.

### Neural activity pre-processing

The spike waveforms of the recorded neuron were isolated and their times were digitized in real time at 1 ms resolution using a two-window spike-amplitude discriminator. For most analyses, the digitized spike times in each trial were converted into a continuous pseudo-analog signal using the partial inter-spike intervals that fell within each sequential time bin in a trial (Cisek and Kalaska, 2005; Coallier et al., 2015; Georgopoulos et al., 1982; Kalaska et al., 1989). Time bin durations varied from 1-20ms in different analyses. Single-trial data were divided into time windows of fixed lengths (e.g. 5ms or 20 ms) or into variable-duration sequential trial epochs for different analyses. An automatic algorithm counted the numbers of whole and fractional inter-spike intervals that fell within the time window or trial epoch. If an inter-spike interval spanned two or more contiguous time windows or epochs, each window or epoch received a fractional count proportional to the fraction of the inter-spike interval that fell within its boundaries. The partial-spike scores were then converted to single-trial spikes/s discharge rates by normalizing for the duration of the time window or trial epoch. Mean cell response histograms were generated by aligning all the single-trial data to different time points in the trial, summing the single-trial discharge rates in corresponding time bins across all trials and then normalizing by the numbers of trials.

All data files were also pre-processed by an automatic algorithm to identify the time of the movement onset (the Reaction Time), and any changes in direction during the reaching movement. The results of this automated analysis of reach kinematics were visually verified for every trial and were corrected manually when necessary (see Coallier et al 2015 for details).

### Quantification and statistical analyses (for both Monkey T and Monkey Z)

Data were analyzed using custom scripts in MATLAB (The Mathworks, Inc) developed and shared by the two labs. Note that both the box-car smoothing and inter-spike interval approaches used in the two labs to convert spike times into firing rates are well-established. All results were fundamentally identical when the two discharge-rate conversion algorithms were applied to the same data files (results not shown).

*i) Psychophysical threshold*

Psychophysical performance was fit to a cumulative Weibull function using the *fit* function of the MATLAB 2018A curve-fitting toolbox. Where *x* is checkerboard coherence, and *p* is the proportion of correct responses:

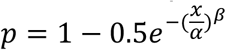

The α parameter is our stimulus threshold, as it is equivalent to the checkerboard coherence at which performance reaches 81.6% correct responses.

*ii) Rapid response changes evoked by the appearance of the first visual cue*

To identify abrupt overt response changes in single-unit activity elicited by first visual cue in each task, we aligned all single-trial data for each unit to the onset of the first cue. Activity was pooled across all trials without regard to checkerboard coherences or eventual reach directions. We then tested the distributions of single-trial activity in each 20 ms bin against the 20 ms bin two bins previously (for instance, the 40-60 ms bin vs the 0-20 ms bin), incremented in 20 ms steps from 0-600 ms after the appearance of the first cue (Wilcoxon signed rank test, p < 0.01). A unit was identified as having a significant abrupt change in activity if it showed a significant change in two consecutive time steps, i.e., significant activity differences spanning an 80 ms time window, and we noted the time bin in which the first significant rapid response change occurred.

*iii) Slope of choice selectivity signal*

To calculate the directional choice selectivity signal (Chandrasekaran et al, 2017; Meister and Huk, 2013), we aligned the single-trial neural activity to the onset of the first and second visual cues in each trial, using both correct and incorrect target-choice trials. We then averaged the single-trial firing rate traces for left/right or PD/oPD reaches separately, for each of the checkerboard coherences. The absolute difference in these left and right averages represents the directional choice selectivity signal for each unit (spikes/s) as it evolves over time. Choice selectivity signals were calculated during a time window from 0-300ms after the first and second visual cues in each task. We then used the MATLAB *fit* function (Matlab 2018A curve-fitting toolbox, The Mathworks Inc.) to estimate the onset time and slope of a linear change in activity after the appearance of each visual cue.

*iv) Repeated-measures ANOVA*

A repeated-measures 3-way ANOVA (IBM SPSS version 24) was performed on the mean single-trial discharge rates recorded in each trial epoch. Main factors were chosen reach direction (“Direction”, D), unsigned checkerboard color coherence independent of dominant color (“Strength”, S), and checkerboard dominant color (“Color”, C). The acceptable significance level was set at p < 0.01 (Bonferroni corrected), and the Greenhouse-Geisser correction was used whenever the sphericity assumption was violated. The ANOVA was done using only trials in which the monkeys chose the correct target, to avoid adding a fourth factor (correct/incorrect choice) to the ANOVA design. The trial epochs included the Center-Hold epoch before the first cues appeared, the Targets-observation epoch before the checkerboard appeared (TF), the Checkerboard-observation epoch before the targets appeared (CF) or the checkerboard was extinguished (CFD), the Checkerboard-RT (TF) and Targets-RT (CF/CFD) epochs from the appearance of the second cue to the onset of movement, and the Movement epoch for the duration movement until the arm reached the target and the Target-hold epoch after target entry to the end of the trial (all tasks).

Unsigned evidence Strength here is a measure of the relative level of the dominant color of the checkerboards without consideration of its actual color or its level of support for a particular target direction, as contrasted with the linear regression analysis (Figure 5). It may also be predictive of the level of confidence that the monkeys could have that their perceptual/motor decision in response to a given checkerboard coherence will be correct, based on accumulated experience with the associated success rates (Figure 1B; Drugowitsch et al., 2014; Zylberberg et al. 2016; Montanède and Kalaska, 2017, SfN Abstract).

*v) Time course of correlations with different task factors*

We assessed to what degree variability in each unit’s activity could be explained by checkerboard parameters and the animal’s choice behavior as a function of time in each trial. We created two matrices of the single-trial firing rates, *y*, of each unit’s neural activity in each task calculated in non-overlapping 20 ms time bins, aligned to either the appearance of the first visual cue or the second visual cue in each trial. We also created a design matrix of predictors, *×*, that included a bias term (all ones) and four task parameter predictors, including the direction of the chosen target independent of its color (e.g., left/oPD reach = −1, right/PD reach = +1), the color of the chosen target independent of its direction (e.g., red/blue = −1, green/yellow = +1), the signed checkerboard color coherence favoring the color of a target independent of its direction (ranging from −100% (red/blue) to +100% (green/yellow); the amount of color evidence for one colored target over the other), and the signed checkerboard coherence strength favoring a direction of target choice independent of its color (from −100% for left/oPD to +100% for right/PD; the amount of color-independent evidence supporting one reach direction over the other). The last predictor requires knowledge of the specific target location-color conjunctions in each trial. Data from trials in which the monkeys chose the correct or incorrect target in each trial were included in this regression analysis so that the color of the chosen target can serve as a surrogate of the monkeys’ sensory interpretation of the color evidence provided by the checkerboard independent of its correct dominant color. Results were similar with and without predictor normalization (e.g., −1 to +1 for all predictors).

For each unit, the firing rate matrix was regressed against the design matrix using the Matlab function *regress* (Matlab 2018A, the Mathworks Inc) to yield predictor weights and confidence intervals for each of those weights at each 20ms time step, using an alpha value of a = 0.001. If the confidence interval for a predictor’s weight did not include 0, then some of the variability in firing rate at this time point was significantly explained by variability in this predictor. For each predictor, we calculated the proportion of units in the population at each 20ms time step for which the predictor’s confidence interval does not include 0. The time series of significant counts for each predictor reflects how the impact of that predictor on single-unit neural activity across the population evolved in time during a trial.

*vi) Ideal-observer analysis of the presence and time course of significant detectability of different task factors in the activity of the neural sample population*

We performed a Receiver Operating Characteristic (ROC) analysis at successive 20ms time intervals to assess the ability of an ideal observer to determine either the dominant color of the checkerboard or the direction of the chosen reaching movement from the distributions of recorded neural activity at different moments in time in a trial, in each task separately. For each unit, data from trials with both correct and incorrect target choices in each task were sorted into two groups according to the dominant color of the checkerboard or the chosen reach direction in each trial pooled across all checkerboard coherence levels. Single-trial firing rates were calculated at 20ms time steps relative to the appearance of either the first or second cue for each of the two groups of trials. The two distributions of firing rates in each time bin were used to calculate the area under the ROC curve (AUC) at that time step for a given unit. This provided a time series of AUC measures for each unit for either the checkerboard color or chosen reach direction. An AUC value of 0.5 indicates an inability to differentiate the two data distributions for either the two colors or the two reach directions, while a value of 1 indicates a perfect ability to distinguish the two. This was repeated for each unit in the sample neural populations in each task (for monkey T, only units tested in both the TF and CFD tasks were used). This yielded distributions of the AUC measures for the sample populations at each 20ms time step for each comparison (checkerboard color or chosen reach direction). To determine whether an ideal observer of the neural activity could show an improvement in their ability to distinguish between the checkerboard colors or chosen reach directions at each time step, we compared the distributions of AUC values in each 20ms bin against the AUC values calculated in a baseline time step −200ms to −180ms before the onset of the first visual cue in each trial (Wilcoxon 1-tailed signed-rank test, p < 0.001).

The ROC analysis was also used to test for a difference in the onset latency of an improvement in the detectability of the chosen reach direction between the two tasks. For monkey T, the AUC values were recalculated at 1ms time steps, starting −200ms before the appearance of the second visual cue, and ending 600ms after its appearance. The population distributions of AUC values at each time step were tested against a baseline time step −200ms before the appearance of the second cue (Wilcoxon 1-tailed signed rank test, p = 0.05/801 = 6.2E-05). The onset latency for each task was identified as the first time step after the appearance of the second visual cue that had a significant increase in AUC values and was followed by 49 successive time steps with a significant increase (i.e., 50ms of uninterrupted significantly larger AUC values compared to the pre-cue baseline activity). The same procedure was used for monkey Z, but at 10ms resolution.

To assess the reliability of that estimate of a difference in onsets, we randomly resampled the unit activity with replacement in each task 1000 times and recalculated the bootstrapped sample onset latencies as just described. We then tested for a significant difference in the distributions of latency estimates (Wilcoxon 1-tailed signed-rank test).

### Data and software availability

Data and software are available upon request.

## Supplemental Material

**Supplemental Figure 1.**
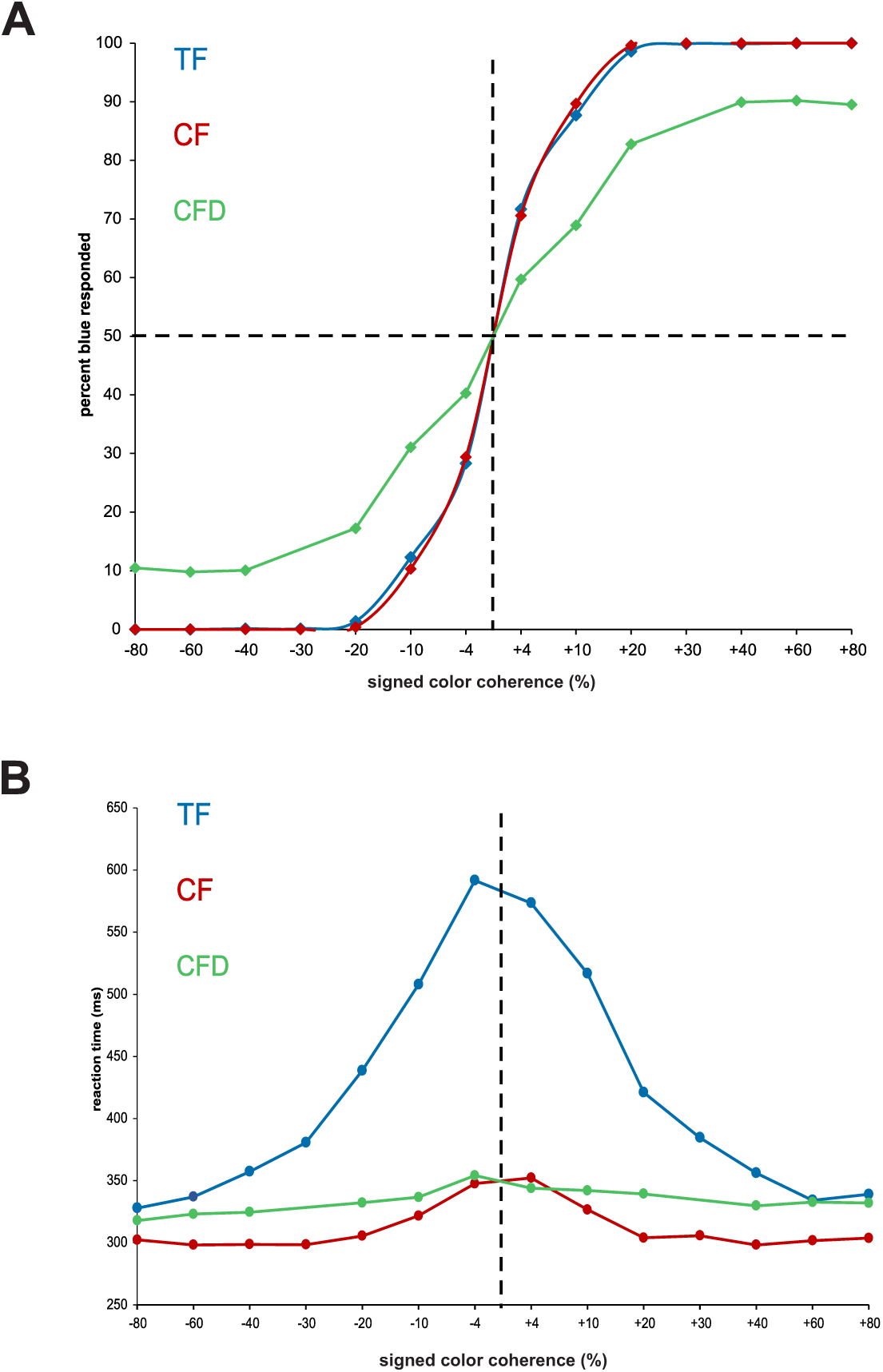
Monkey Z behavior for all tasks: TF, CF, and CFD. **A)** Psychometric curves and B) chronometric curves are shown; compare to Figure 1 **B,C**). TF and CF tasks: same data as in Figure 1B,C. Like monkey T, monkey Z showed a decrease in success rates for the checkerboards with the highest color coherences in the CFD task compared to the TF and CF tasks. Unlike monkey T, however, monkey Z continued to show a substantial reduction in RTs for checkerboards with lower color coherences in the CFD task compared to the TF task.

**Supplemental Figure 2.**
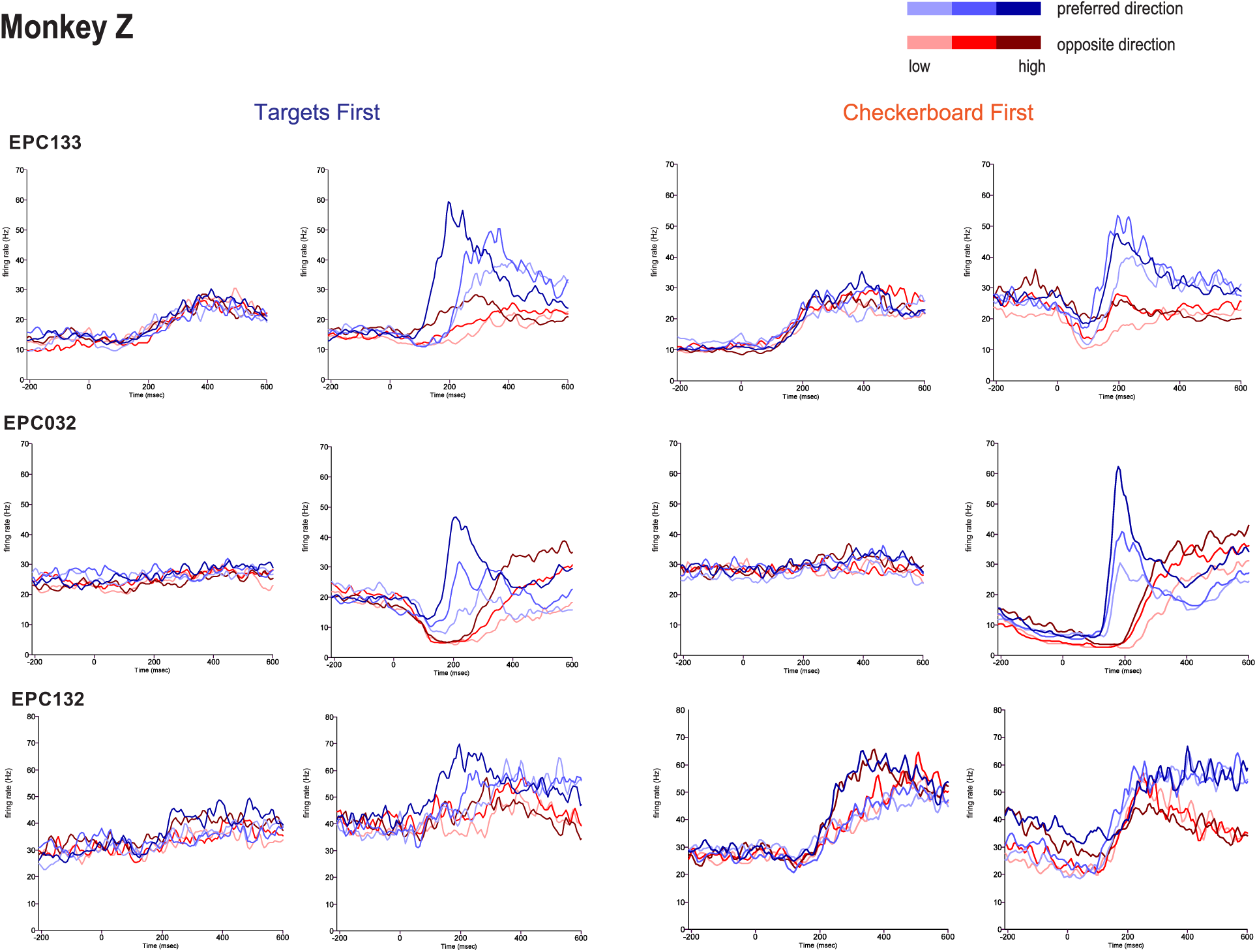
Further examples of units with response to first visual cue onset; same format as Figure 3. Unit EPC133 showed a significant rapid response onset (see Methods) at 220ms after the Targets cue appeared in the TF task and 180ms after the Checkerboard appeared in the CF task. Unit EPC032 did not show a significant rapid response to the Targets in the TF task but showed a late rapid decrease in activity (>600ms) to the Checkerboard in the CF task. Unit EPC132 showed a significant rapid increase in response to the Targets at 280ms after they appeared in the TF task, and a significant rapid response increase 220ms after the Checkerboard appeared in the CF task. The response to the Checkerboard was markedly stronger and more rapid for the 100% coherence than for the 60% and 52% coherences. EPC132 showed a significant main effect of evidence Strength in the ANOVA, but did not show a significant linear regression to either the signed color coherence or the signed evidence for reach direction during the Checkerboard-observation period of the CF task.

**Supplemental Figure 3.**
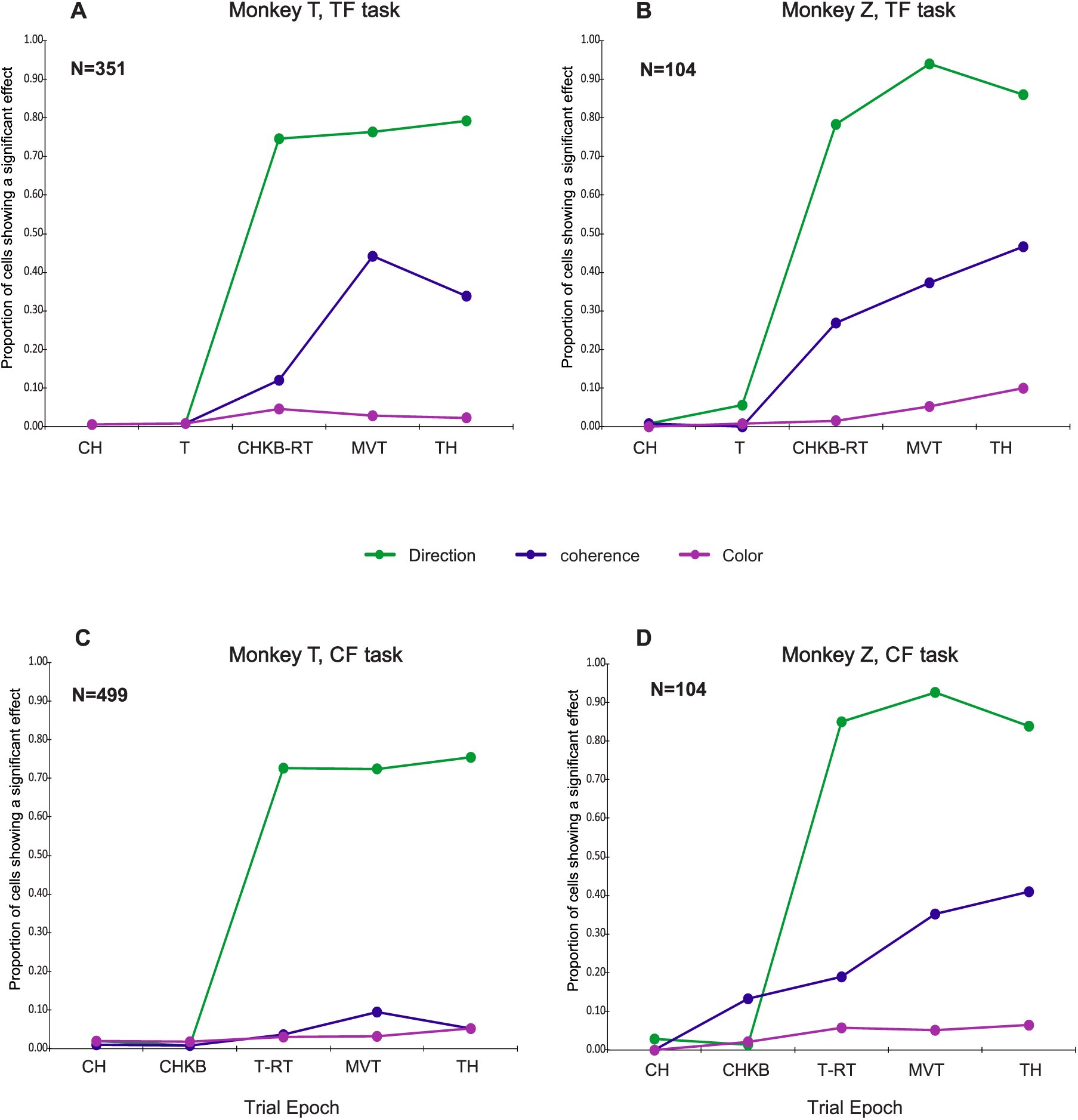
Proportion of the units that showed a significant main effect of the task factors chosen reach Direction, unsigned checkerboard evidence Strength and checkerboard dominant Color in different trial epochs (ANOVA, p< 0.01). **A,B**) TF task. Very few neurons showed a significant main effect of any task factors during the Center-hold (CH) and Targets-observation (T) epochs. Furthermore, very few units in either monkey showed a significant modulation by the color-location conjunction information provided by the Targets cue during the Targets-observation period (Direction × Color interaction, p < 0.01; Supplemental Table 1). After the checkerboard appeared, many units in both monkeys showed a significant effect of Direction (green) during the Checkerboard-RT epoch before the onset of the reaching movements (CHKB-RT), during the Movement epoch toward the targets (MVT) and during the Target-Hold epoch at the end of the reaching movements (TH). A smaller proportion of units showed a significant main effect of checkerboard evidence Strength (blue) during the CHKB-RT, MVT and TH epochs. Very few units in either monkey showed a significant effect of checkerboard dominant Color (magenta) in any trial epoch. Many neurons in both monkeys also showed a significant effect of evidence Strength on their Direction responses (significant Direction × Strength interaction, p < 0.01). The effects of evidence Strength and the Direction × Strength interaction captured most of the effect of checkerboard coherence on the directional activity of the neurons in the TF task in all trial epochs after the checkerboard appeared (Figure 2, 3; Supplemental Figure 2). **C,D**) CFD/CF task. Very few units in either monkey showed a significant main effect of Direction during the CH and Checkerboard-observation (CHKB) epochs. Many units emitted strongly Direction signals during the Targets-RT (T-RT), MVT and TH epochs. In monkey T (**C**), relatively few units showed a significant main effect of checkerboard evidence Strength. In contrast, some units in monkey Z (**D**) were significantly modulated by checkerboard Strength during the CHBK epoch, and the number of significant effects of Strength increased progressively during the T-RT, MVT and TH epochs, but fewer than in the TF task. Very few units in either monkey showed a significant effect of checkerboard dominant Color, or Color × Direction or Color × Strength interactions in any trial epoch (Supplemental Figure 4; Supplemental Table 1).

**Supplemental Table 1.** ANOVA results for monkey T and monkey Z; see Supplemental Figure 4. D, chosen reach direction; C, checkerboard dominant color; S, unsigned checkerboard coherence strength; DxC, DxS, CxS, interactions between variables.

| Targets First Task |  |  |  |  |  |  |  |  |  |  |  |  |  |  |  |  |  |  |  |  |  |  |  |  |  |  |  |  |  |  |  |  |
| --- | --- | --- | --- | --- | --- | --- | --- | --- | --- | --- | --- | --- | --- | --- | --- | --- | --- | --- | --- | --- | --- | --- | --- | --- | --- | --- | --- | --- | --- | --- | --- | --- |
| Epoch |  | Center-hold |  |  |  |  |  | Targets |  |  |  |  |  | Checkerboard-RT |  |  |  |  |  | Movement |  |  |  |  |  | Target-hold |  |  |  |  |  |  |
| Factor |  | D | C | S | DxC | DxS | CxS | D | C | S | DxC | DxS | CxS | D | C | S | DxC | DxS | CxS | D | C | S | DxC | DxS | CxS | D | C | S | DxC | DxS | CxS |  |
| Monkey T | Units | 2 | 2 | 2 | 8 | 2 | 3 | 3 | 3 | 3 | 1 | 2 | 0 | 260 | 16 | 42 | 7 | 16 | 2 | 266 | 10 | 154 | 6 | 61 | 5 | 276 | 8 | 118 | 3 | 57 | 1 | 3 |
|  | Decimal fraction | 0.01 | 0.01 | 0.01 | 0.02 | 0.01 | 0.01 | 0.01 | 0.01 | 0.01 | 0.00 | 0.01 | 0.00 | 0.74 | 0.05 | 0.12 | 0.02 | 0.05 | 0.01 | 0.76 | 0.03 | 0.44 | 0.02 | 0.17 | 0.01 | 0.79 | 0.02 | 0.34 | 0.01 | 0.16 | 0.00 | 5 |
| Monkey Z | Units | 1 | 0 | 1 | 0 | 0 | 2 | 5 | 1 | 0 | 5 | 1 | 1 | 79 | 2 | 26 | 5 | 10 | 1 | 95 | 7 | 39 | 4 | 19 | 2 | 86 | 10 | 49 | 2 | 20 | 2 | 1 |
|  | Decimal fraction | 0.01 | 0.00 | 0.01 | 0.00 | 0.00 | 0.03 | 0.06 | 0.01 | 0.00 | 0.05 | 0.01 | 0.01 | 0.78 | 0.01 | 0.27 | 0.04 | 0.09 | 0.01 | 0.93 | 0.05 | 0.37 | 0.04 | 0.18 | 0.01 | 0.85 | 0.10 | 0.46 | 0.02 | 0.20 | 0.01 | 1 |

| Checkerboard First with Delay / Checkerboard First Task |  |  |  |  |  |  |  |  |  |  |  |  |  |  |  |  |  |  |  |  |  |  |  |  |  |  |  |  |  |  |  |  |
| --- | --- | --- | --- | --- | --- | --- | --- | --- | --- | --- | --- | --- | --- | --- | --- | --- | --- | --- | --- | --- | --- | --- | --- | --- | --- | --- | --- | --- | --- | --- | --- | --- |
| Epoch |  | Center-hold |  |  |  |  |  | Checkerboard |  |  |  |  |  | Targets-RT |  |  |  |  |  | Movement |  |  |  |  |  | Target-hold |  |  |  |  |  | M |
| Factor |  | D | C | S | DxC | DxS | CxS | D | C | S | DxC | DxS | CxS | D | C | S | DxC | DxS | CxS | D | C | S | DxC | DxS | CxS | D | C | S | DxC | DxS | CxS |  |
| Monkey T | Units | 10 | 10 | 5 | 1 | 5 | 3 | 4 | 9 | 4 | 4 | 1 | 5 | 360 | 15 | 18 | 17 | 3 | 1 | 359 | 16 | 47 | 5 | 15 | 2 | 374 | 26 | 26 | 14 | 17 | 4 | 45 |
|  | Decimal fraction | 0.02 | 0.02 | 0.01 | 0.00 | 0.01 | 0.01 | 0.01 | 0.02 | 0.01 | 0.01 | 0.00 | 0.01 | 0.72 | 0.03 | 0.04 | 0.03 | 0.01 | 0.00 | 0.72 | 0.03 | 0.09 | 0.01 | 0.03 | 0.00 | 0.75 | 0.05 | 0.05 | 0.03 | 0.03 | 0.01 | 45 |
| Monkey Z | Units | 3 | 0 | 0 | 0 | 2 | 2 | 1 | 2 | 12 | 0 | 1 | 0 | 88 | 6 | 22 | 4 | 8 | 0 | 94 | 6 | 38 | 4 | 18 | 1 | 84 | 7 | 44 | 5 | 15 | 2 | 10 |
|  | Decimal fraction | 0.03 | 0.00 | 0.00 | 0.00 | 0.02 | 0.02 | 0.01 | 0.02 | 0.13 | 0.00 | 0.01 | 0.00 | 0.84 | 0.06 | 0.19 | 0.04 | 0.07 | 0.00 | 0.92 | 0.05 | 0.35 | 0.03 | 0.16 | 0.01 | 0.83 | 0.06 | 0.41 | 0.05 | 0.16 | 0.02 | 10 |

| Table Legend |  | 1231 |
| --- | --- | --- |
| D: Direction chosen |  | DxC: interaction between Direction and Checkerboard dominant color 1232 |
| C: Checkerboard dominant color |  | DxS: interaction between Direction and Checkerboard strength 1234 |
| S: Checkerboard strength |  | CxS: interaction between Checkerboard dominant color and Checkerboard strength 1236 |

### PMd neurons in monkey Z but not monkey T exhibited responses to the first sensory information provided in each task

The response of the PMd neurons in monkey Z to the appearance of the Target cues in the TF task is consistent with a similar activation of PMd neurons in a 2-Target task, and has been interpreted as a simultaneous co-activation of two PMd populations that prefer reaches to the two potential targets before a second monochromatic instructional cue identifies the correct target (Cisek and Kalaska, 2005; Coallier et al., 2015). This activity during the memory delay period of the 2-Target task is a neural correlate of a memorized representation of the Target cue information in each trial, expressed in the space of potential actions in PMd.

The responses of the neurons to the appearance of the Checkerboard cues in the CF task likely do not implicate PMd in the perceptual interpretation of the color evidence since the responses were largely insensitive to the salient dimensions (dominant color, signed color coherence) of the stimuli (Figure 5, 6). Alternatively, the activation may reflect a similar process of representation of potential actions as in the TF task. Once a neuron in monkey Z was chosen for recording in a given recording session, it was tested with the same two opposite target locations in the task-defined preferred reach direction of the neuron and the opposite direction for hundreds of trials in the two tasks. Therefore, it is reasonable to expect that monkey Z had some level of implicit prior knowledge of where the two targets would appear at the start of each trial in both tasks and all that it lacked was the specific color-location conjunction, which it obtained when the targets appeared. The activity during the Checkerboard observation period in the CF task may have reflected the accumulating knowledge of the dominant color of the checkerboard in neural circuits outside of PMd. This may have enabled a covert activation of the simultaneous representations of the two anticipated potential actions even before the targets actually appeared in the CF task since that accumulating sensory evidence will eventually support one of the colored targets once they appeared. This coactivation of the two PMd populations preferring the two targets may have in turn contributed to the shorter RTs in the CF task than the TF task for even the 100% checkerboards. Critically, however, the linear regression (Figure 5), ROC (Figure 6) and ANOVA (Supplemental Figure 3) showed that these activations during the first-cue observation periods in PMd of monkey Z did not reflect the final decision-related processes as defined here because it did not predict any aspect of monkey Z’s differential choice behavior after the second visual cue appeared in each task.

Since activation of PMd by partial action-related information before the final action choice can be specified is a robust finding in several different studies, why did monkey T not show any activity in PMd during the first visual-cue observation periods in the TF and CFD tasks? This could occur if different cortical regions were sampled in monkeys T and Z. Both monkeys are still participating in neural recordings and histological localization of penetration sites has not yet been done. However, similar stereotactic coordinates for chamber implants and extensive relative overlap of recording penetrations within the chambers in the two monkeys suggest that is not the main cause (Figure 1E).

Another possible explanation is the different training history of the two monkeys. Monkey T only ever experienced L/R targets and was initially trained in the TF task in which the targets remained visible for the duration of each trial. Thus, its original training experience did not require the establishment of a memorized trace of the target information. In contrast, monkey Z was trained for many months in the 1T and 2T tasks with targets in 8 different directions that varied from trial to trial and with two long memory-delay periods during which the monkey had to remember the spatial location (1T task) and color-location conjunctions of the targets before selecting a target (2T task), and initiating a reach (1T and 2T tasks). The 1T and 2T tasks were also used in all neural recording sessions to search for task-related neurons. Engagement of PMd during the memory-delay periods of the 1T and 2T tasks may have facilitated task performance for monkey Z, which was carried over to the TF and CF tasks.

Another possible contributing factor is the target location placement in the tasks. For monkey Z, targets were placed in spatial locations along their preferred-opposite movement direction axis to maximize the directional activity difference for each unit in the TF and CF tasks, and would change from neuron to neuron. For monkey T, in contrast, target locations were fixed to the left and right of center, and units were recorded for that single movement axis, regardless of what their preferred reach directions might have been.

Finally, the lack versus presence of activity changes after the checkerboards appeared in the CF/CFD tasks may have reflected a difference in the strategy that the two monkeys adopted to perform the tasks. Monkey T may have largely deferred the interpretation of the checkerboard evidence until the appearance of the targets, resulting in no PMd responses and longer RTs in the CFD task compared to the TF task. In contrast, monkey Z appeared to largely commit to a categorical decision about the dominant color of the checkerboard while observing it in the CF task, resulting in a substantial reduction in RTs after the targets appeared. This may have been accompanied by a covert activation of the two competing action-related PMd populations like in the TF task, even though the targets had not yet appeared. This is further reinforced by the finding that monkey Z’s RTs remained drastically shorter in a modified version of the CF task that had the same temporal structure as the CFD task. This showed that monkey T’s prolonged RTs in the CFD task compared to the TF task were not due to the imposed memory-delay period but rather to how and when it interpreted that task-relevant sensory evidence provided by the checkerboards.

### How did the monkeys convert checkerboard color coherence into reach actions?

The systematic differences in the rate of rise of reach-related directional signals in PMd as a function of the color coherence of the checkerboards in the TF task are reminiscent of similar correlations with the coherent motion strength of RDK stimuli seen in saccade-related cortical regions (Kim and Shadlen, 1999; Shadlen and Newsome, 2001; Roitman and Shadlen, 2002) and in parietal cortex area 5 in an arm reach task (de Lafuente and Shadlen, 2015). Those findings have been interpreted as the neural correlate of a process of accumulation of noisy sensory evidence across time using signals generated by motion-sensitive neurons in medial temporal cortex (MT) to inform the choice of the action that must be performed to report a decision about perceived net motion direction. This makes intuitive sense since the RDK stimuli are stochastic, contain variable amounts of dot motion in random directions as well as in the coherent motion direction from moment to moment, and only evoke reliable sensations of coherent visual motion in a particular direction when experienced over time (Roitman and Shadlen, 2002).

In contrast, the checkerboards used in these tasks comprised sets of small squares whose colors are easily and rapidly discriminable, and contained no input signal “noise” comparable to the variable numbers of dots moving in the coherent and random directions from frame to frame in the RDK stimuli. The checkerboard stimuli presented in half of the trials to monkey Z were dynamic and changed every 50ms, but the numbers of blue and yellow squares in each stimulus stream remained fixed in a given trial and only their positions within the checkerboard changed from image frame to image frame. In contrast, in the other half of the trials for monkey Z and all of the trials for monkey T, a single static checkerboard of R/G or B/Y squares appeared for the duration of the observation period in each trial, so that the physical properties of the sensory stimulus that the monkeys experienced did not change across time. Despite these differences in the visual stimuli, the two monkeys showed remarkably similar chronometric and psychophysical trends in the TF task (present study; Chandrasekaran et al., 2017). Furthermore, human and non-human subjects also showed very similar performance when viewing either dynamic or static checkerboard stimuli (Coallier and Kalaska 2014; Coallier et al 2015; present study). Thus, whereas the motion sensations evoked by RDK stimuli require dynamically changing stimuli across time, the assessment of the color evidence in the checkerboards was relatively insensitive to the presence or absence of continually updated sensory inputs. This does not, however, preclude momentary stochastic noise generated within the central neural circuits that process even the static checkerboard visual input (both monkeys) and store it in short-term working memory (monkey T).

The similarity of task performance in the TF task is also striking given another difference in the checkerboard stimuli experienced by the two monkeys. The checkerboards used in monkey T’s experiments contained only task-salient R and G squares. In contrast, the checkerboards used in monkey Z’s experiments contained 100 task-salient B and Y squares against a background of 125 task-irrelevant R squares. This reduced the overall density of task-relevant color information in the checkerboards for monkey Z and required it to identify the task-relevant information from among the “distractor” red squares. Despite this difference, the psychophysical curves and psychophysical thresholds of the two monkeys were very similar in the TF task, and the RTs for the high-coherence checkerboards were actually shorter in monkey Z than monkey T (Figure 1).

Despite the absence of color evidence “noise” in the checkerboards and the insensitivity of task performance to static versus dynamic stimuli, the monkeys took longer to choose a colored target when the checkerboard coherence decreased (Coallier and Kalaska 2014; Coallier et al 2015; Chandrasekaran et al., 2017). In RDK stimuli, this effect has been explained by a sequential-sampling process that takes longer to identify the direction of the weak coherent-motion signal generated by MT neurons against a high level of motion direction noise. For the checkerboard stimuli, in contrast, it presumably reflects a longer period of time required to determine whether the checkerboard was predominantly one or the other of the two task-salient colors as the numbers of squares of the two easily-discriminable colors became more similar. This may require longer re-sampling of the sensory input while observing the checkerboard (monkey Z) or from a noisy working-memory trace of the checkerboard (monkey T; Pearson et al., 2014; Shadlen and Shohamy, 2016, Shushruth and Shadlen, CoSyNe abstract). We can assume that the color evidence is initially processed by neurons in the parvocellular “color-opponent” pathway (Conway and Livingstone, 2006; Conway, 2014; Bohon et al., 2016; Cheadle and Zeki, 2014). However, to our knowledge, there have been no studies of neural responses in that pathway to multi-colored checkerboard stimuli like ours in color discrimination tasks analogous to the many studies of visual motion processing in MT.

The perceptual decision could be considered as a pure color discrimination problem since the subjects had to estimate the dominant color of the checkerboards in order to identify the reach target whose color matched that of the checkerboard. However, similar dichromatic dot arrays have been used in studies of numerosity, the ability of subjects to estimate relative numbers of visual objects (Cantlon et al., 2009; Ratcliff and McKoon, 2018; Burr et al., 2017). Subjects likewise showed longer RTs when the relative numbers of objects in the stimuli are similar (Ratcliff et al., 2015; Ratcliff and McKoon, 2018). These results have been interpreted as consistent with a process of sequential sampling and accumulation of evidence across time and across space within the stimuli (Ratcliff, 2014; Ratcliff et al., 2015; Ratcliff and McKoon, 2018; Fornaciai and Park, 2017), but did not speculate on the nature of the sensory evidence that was being sampled, unlike RDK stimuli. Furthermore, the checkerboards that we used have inherent in them several potential confounding “low level” physical properties identified in numerosity studies that are independent of the presumably “higher level” sense of relative numbers per se, including the relative area of the checkerboard occupied by squares of each color, their total circumference, and the relative degree of spatial contiguity of squares of the same color (i.e., how often they cluster to share a common border) (Gebuis and Reynvoet, 2012; Leibovich and Henik, 2013; Dietrich et al., 2015; Ratcliff and McKoon, 2018; Harvey and Dumoulin, 2017). Indeed, the colored squares did not have a neutral-colored border and so would form larger monochromatic “clumps” when contiguous, rather than remaining visible as discrete squares (Figure 1D). All of these factors could have contributed to the monkeys’ estimation of the dominant color of the checkerboards, independent of any estimate of relative numbers of squares. Furthermore, the number and density of squares in our checkerboards were usually higher than normally used in numerosity studies and more closely resemble what are called “textures”, which follow different psychophysical laws than dot arrays with smaller numbers of elements (Burr et al., 2017).

However, this study was not designed to study numerosity or to examine what specific properties of the checkerboards the subjects used to make the relative color estimates. Instead, the checkerboards were chosen as a means to present stimuli with different levels of competing evidence for two alternative reach choices, using a stimulus dimension (color) that has no inherent natural association with the directionality of motor output. Our findings indicate that PMd neurons process information pertaining to the likelihood of different action choices provided by the checkerboard stimuli, independent of the critical decision-relevant physical property of the sensory input on which those action likelihoods are based, in this case its dominant color.

Important questions not directly addressed by this study are where are the neural correlates of the critical color-related information on which the action decisions were based and how are they transformed into color-independent evidence supporting the action choices? A strong candidate is the dorsolateral prefrontal cortex (Di Pellegrino and Wise, 1991; Mante et al., 2013). We have preliminary evidence that the specific color/location conjunctions of the spatial target cues and color/location matching rules after the checkerboard appeared in each trial are expressed in lateral prefrontal cortex around the principal sulcus while a monkey performed a TF task (Coallier et al., 2008, SfN abstract).

The effect of checkerboard color coherence on task performance and neural activity is consistent with a number of different computational decision-making models, including drift-diffusion (Gold and Shadlen, 2007; Ratcliff et al., 2016; Ratcliff and McKoon, 2018; Roitman and Shadlen 2002; Shadlen and Newsome 2001); gated stochastic accumulation (Schall et al., 2011; Purcell et al., 2012), urgency gating (Cisek et al 2009; Thura and Cisek 2014), and independent-race (Carpenter and Williams, 1995; Noorani and Carpenter, 2016; Brown and Heathcote, 2008). Nevertheless, until more neurophysiological findings are available about the sources and nature of sensory signals that are being processed while subjects estimate the relative amounts of colored squares in the dichromatic checkerboard stimuli, and how those sensory signals are transformed into action-related information, we prefer to remain agnostic as to the computational mechanisms that underlie the task performance of the subjects.

